# A Minimal Genetic Circuit for Cellular Anticipation

**DOI:** 10.1101/2025.04.22.649979

**Authors:** Jordi Pla-Mauri, Ricard Solé

**Affiliations:** ICREA-Complex Systems Lab, Universitat Pompeu Fabra, Dr. Aiguader 88, 08003 Barcelona; Institut de Biologia Evolutiva, CSIC-UPF, Pg. Maritim de la Barceloneta 37, 08003 Barcelona; Santa Fe Institute, 1399 Hyde Park Road, Santa Fe NM 87501, United States

**Keywords:** Anticipation, Basal cognition, Stochastic environment, Adaptive behavior

## Abstract

Living systems have evolved cognitive complexity to reduce environmental uncertainty, enabling them to predict and prepare for future conditions. Anticipation, distinct from simple prediction, involves active adaptation before an event occurs and is a key feature of both neural and aneural biological agents. Building on the moving average convergence–divergence principle from financial trend analysis, we propose an implementation of anticipation through synthetic biology by designing and evaluating experimentally testable minimal genetic circuits capable of anticipating environmental trends. Through deterministic and stochastic analyses, we demonstrate that these motifs achieve robust anticipatory responses under a wide range of conditions. Our findings suggest that simple genetic circuits could be naturally exploited by cells to prepare for future events, providing a foundation for engineering predictive biological systems.

## I. INTRODUCTION

Living agents evolved cognitive complexity as a way to reduce environmental uncertainty or, in other words, as a way to predict their worlds better [1–4]. Anticipatory behavior is a key component of cognition in neural agents, i.e., multicellular organisms equipped with neuron-based circuits and sensors, but not limited to them [5–9]. In addition to memory and learning, anticipation is an essential component of the toolkit for basal cognition [10], has played a crucial role in the evolution of cognitive agents [11], and can play a key role in organismal homeostasis [12]. Theoretical work on anticipation traces back to cybernetics [13, 14] and was formalized by Robert Rosen, who described it as a system’s capacity to prepare for future events using available knowledge [15]. Unlike simple prediction, anticipation involves an active process in which a system adapts and responds before an event occurs [16, 17].

Rosen distinguished three components required to build an anticipatory system, sketched in Figure 1a. First, the system Σ is understood as some cognitive agent (but can also be mechanical) interacting with its environment. A system processes information and makes adjustments based on its anticipated future states. Secondly, within this system, a model *M* (a representation of reality) enables the system to predict outcomes. Finally, anticipation involves the controller *C* (also known as the effector), which is the component that transforms anticipation into action. It allows the system to respond proactively rather than reactively. When the model predicts a future event, the effector allows the system to adjust its behavior in advance [15].

**Figure 1.**
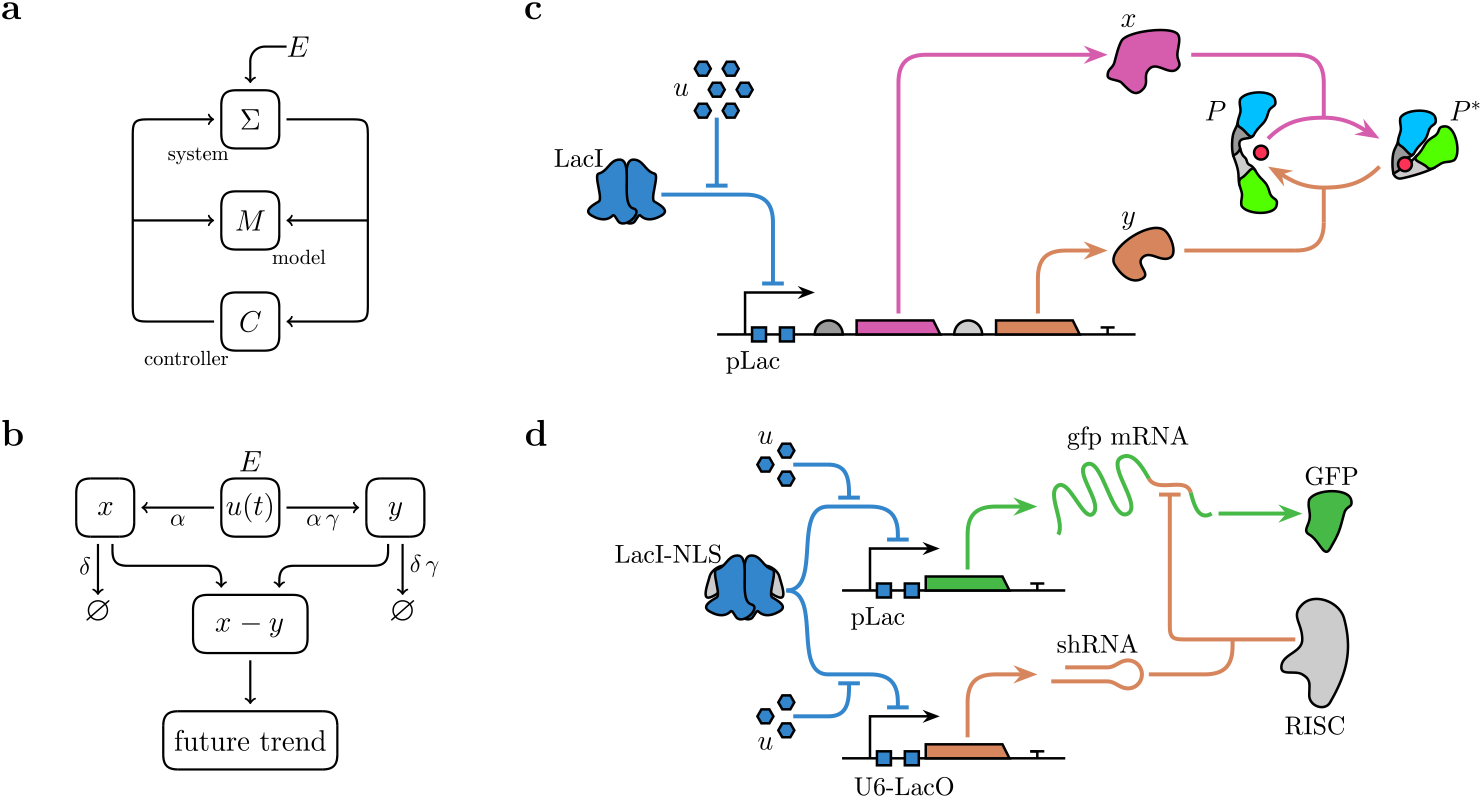
Anticipatory circuits. In (a), a basic diagram of Rosen’s anticipatory system is shown, involving three key components. In Frank’s model (b), anticipation is achieved by dealing with trends using the difference *x* − *y* between two signals displaying different timescales of response. In (c), a genetic circuit is designed to predict the incoming signal *u*; which in this case is either IPTG or lactose, an external inducer of the *lac* promoter (pLac) under LacI repression. Upon binding, *u* triggers the concurrent expression of both kinase *x* and phosphatase *y* from a single operon. The relative expression levels of these proteins are controlled by varying the sequences upstream of each coding region—such as ribosome binding sites in prokaryotes or Kozak sequences in eukaryotes. Additionally, the kinase can be fused with a degradation tag to increase its turnover rate. The dynamic kinase : phosphatase activity ratio can then be monitored in real time using a FRET-based sensor, with a ligand targeted by both enzymes. An alternative design is illustrated in (d), where the input signal *u* triggers the co-expression of a shRNA and mRNA containing a degradation target. The shRNA-derived complex reduces the amount of translatable mRNA through active degradation at the 3^′^ UTR region, leading to decreased production of fluorescent protein. As a result, an increase in fluorescent response will be detected when a positive trend in the input is anticipated.

Anticipation is also present in single cells, providing a strong adaptive advantage by allowing faster responses to changing conditions, which can improve survival [10, 18]. An example is provided by chemotaxis, i.e. the movement of bacteria towards beneficial chemicals, like nutrients, and away from harmful ones by sensing and responding to chemical gradients. This behavior is anticipatory because bacteria do not simply react to stimuli but compare past and present conditions to predict future outcomes. Adjusting their movement according to environmental trends, they effectively “anticipate” better conditions, demonstrating a basic form of forward-looking behavior [19, 20]. Oscillatory genetic circuits can also contribute to anticipatory behavior, especially in cells that experience periodic environmental changes. Circadian rhythms in cyanobacteria, for example, allow them to anticipate daily light–dark cycles and optimize photosynthesis [21, 22].

Some simple genetic motifs can be used to design genetic circuits that implement this feature. Minimal genetic circuits enabling anticipation can be obtained from the feedforward loop (FFL), where an input regulates an intermediate factor and a final output. In an incoherent type-1 FFL, for instance, the input activates both the target gene and a repressor, creating a transient anticipatory response [23]. Empirical evidence suggests that simple cellular organisms could have evolved to discern and use temporal correlations in environmental signals to predict future conditions [24]. A notable example of this is sugar metabolism within the intestinal tract, where the detection of one sugar prepares the cell for the subsequent encounter with another [5, 25]. A similar adaptive response has also been achieved under experimental evolutionary conditions [26]. There is thus a diversity of possible designs, but what is the minimal implementation allowing for anticipation?

In a recent paper, Steven Frank proposed a simple design principle for a biological system capable of anticipating trends [27]. This circuit (Figure 1b) computes the difference between short- and long-term moving averages, analogous to financial moving average convergence– divergence indicators commonly used in financial markets to predict asset price trends. This architecture effectively tackles core signal processing challenges: for noisy inputs, moving averages suppress noise through temporal integration, yielding a more accurate estimate of the current state; for non-Markovian signals—where future behavior depends on historical states rather than just the present—the differential output captures persistent trends by leveraging these existing correlations [19, 20]. As a result, the circuit can estimate the momentum of the current trend and, consequently, predict the future direction of change in the environment. Natural systems employ similar strategies, such as the bacterial *mazEF* toxin–antitoxin system [28]. This system maintains a balance between toxin and antitoxin through differential degradation and stoichiometry, allowing rapid activation of toxin under stress by disrupting this balance and leading to mRNA cleavage.

Within cognitive science, the concept of anticipation (or prediction) has generally been accepted without much controversy. Neural agents—including humans—use their cognitive machinery to make predictions [29–33]. However, the idea that such predictive capacity can also present within aneural systems has been oftentimes controversial [34]. Recent work on learning in aneural systems, however, provides empirical and theoretical support for this possibility [35, 36]. Synthetic biology provides a different way to study the requirements for anticipation and thus gives us another path for exploration. Since engineered circuits allow for full control over the requirements and dynamical outcomes, quantitative measures of anticipation become possible without ambiguities. Previous studies have shown how other key features of basal cognition can be engineered in cells or cellular consortia. These include memory circuits [37, 38], associative learning [39, 40], or collective intelligence [41], but no anticipatory designs have been proposed so far.

Following Frank’s proposal, here we suggest a potential implementation, using synthetic biology, of his basic design principle. A minimal anticipation motif is proposed, and its potential for anticipation is evaluated. The response of this anticipation motif is analyzed for both deterministic (constant, periodic) changes and stochastic fluctuations modeled using Wiener-derived processes [27]. Anticipatory responses are shown to be achieved under a broad range of conditions, suggesting that such minimal circuits could be easily exploited by cells to deal with future events.

## II. METHODS

Our proposed minimal motif can be implemented as a simple two-component genetic circuit, with both components activated by an input signal, *u*. This circuit allows for the accumulation of two molecular species, *x* and *y* at different rates, effectively creating moving averages with distinct timescales. By comparing these averages, the system can effectively determine in which direction the trend is heading [27]. Such moving averages can be modeled by a system of two differential equations,

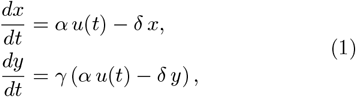

where *α* ≥ 0 is a production rate in response to the input signal, *δ* ≥ 0 represents the degradation/dilution rate, and 0 < *γ* < 1 is a scaling factor that makes one gene respond more slowly than the other, creating the different timescales.

A possible synthetic biological implementation using a push–pull motif can be achieved by assigning kinase activity to *x* and phosphatase activity to *y*, both targeting the same constitutively expressed protein *P* (see Figure 1c). In this configuration, *x* catalyzes the phosphorylation of protein *P* to its active form *P* ^∗^, while *y* catalyzes the dephosphorylation reaction returning *P* ^∗^ to its dephosphorylated state *P* . The circuit output is a constitutively expressed tandem of fluorescent proteins linked by a phosphorylation target sequence. Kinase activity induces phosphorylation-dependent conformational changes that enable Förster resonance energy transfer (FRET) between the fluorophores, while concurrent phosphatase activity reverses this process, thereby modulating FRET efficiency. The dynamic equilibrium between these opposing activities results in measurable spectral shifts in fluorescence emission, which correlate with the kinase:phosphatase activity ratio over time. This ratiometric fluorescence response provides a direct and quantitative readout of the system’s dynamical state in response to input *u*, enabling experimental validation of the proposed theoretical model.

Assuming a normalized steady-state concentration for the total observable *P* + *P* ^∗^ = 1, the dynamics are described by a coupled differential equation system,

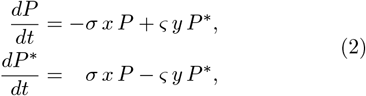

where *σ* and *ς* represent the kinetic rates of phosphorylation and dephosphorylation, respectively. Using the constraint *P* = 1 − *P* ^∗^, the previous system can be reduced to a single equation,

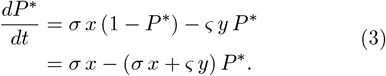

Under the assumption that post-transcriptional modifications occur at a significantly faster timescale than protein production and degradation, *P* ^∗^ is considered to reach quasi-steady-state rapidly, yielding

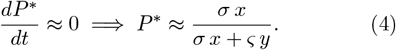

Deviations from this equilibrium indicate the direction of the trend: a higher proportion of active proteins suggests an upward trend, while a lower proportion suggests a downward trend.

An alternative implementation uses direct inhibition of the output, analogous to the synthetic system shown in Figure 1d or natural toxin-antitoxin mechanisms. In the synthetic system, both a short hairpin RNA (shRNA, *y*) and a messenger RNA (mRNA, *x*) containing a homologous sequence are repressed by a shared transcription factor regulated by the input. Once processed and incorporated into the RISC complex, the shRNA guides the sequence-specific cleavage of the mRNA’s 3^′^ untranslated region (UTR), reducing the pool of intact mRNA available for translation into the output fluorescent protein. Consequently, the fluorescent signal accumulates when an increase in the input is predicted, and diminishes otherwise.

Both implementations share a common expression for the point at which a change in trend sign is predicted:

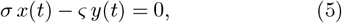

which depends on the kinetic rates and the stoichiometry of the reactions involving *x* and *y*. For simpler analytical tractability, the equivalence *σ* = *ς* will be assumed in all subsequent derivations.

In the next section, several different case studies are considered, dealing with a diverse range of input signals (constant, periodic, fluctuating) and the responses associated with the minimal anticipation circuit presented. As it will be shown, anticipatory behavior is obtained under well-defined ranges of conditions.

## III. RESULTS

### A. Deterministic Input

To explore the anticipatory capabilities of the proposed genetic circuits, we first analyze their response to deterministic input signals. By considering inputs with well-defined dynamics, we can derive analytical solutions that reveal the core principles underlying anticipation.

Most results are derived from an analysis of the long-term dynamics, with full derivations in the Supplementary Material (SM).

#### 1. Constant Drift Input

The simplest case study of a time-dependent input signal would correspond to linear changes. Let us consider first an input signal *u*(*t*) with a constant rate of change, as described by

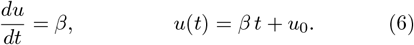

The system’s behavior at long times (*t* → ∞) approaches the following asymptotic forms (see SM I B for full derivation):

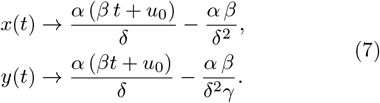

The anticipated trend, determined by the sign of the difference of the moving averages, is then given by

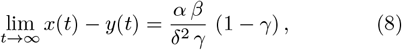

which correctly predicts the trend, as its sign matches that of the input’s slope, *β*.

A more complex scenario involves an input signal whose rate of change is piecewise constant. The input initially changes at a constant rate *β*_1_ during the time interval *t* ∈ [*t*_0_, *t*_*s*_). At time *t*_*s*_, the rate of change abruptly transitions to a new constant value, *β*_2_. This behavior is captured by the following piecewise input function:

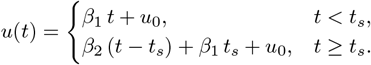

For *t* < *t*_*s*_, the sign prediction is identical to the constant rate input case, as described by equation (8). However, for *t* ≥ *t*_*s*_, assuming *t*_*s*_ sufficiently large such that the transient dynamics before the switch are negligible, and taking *τ* := *t* − *t*_*s*_, the difference of moving averages can be approximated as (see SM I C for full derivation):

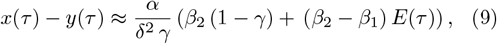

where *E*(*τ*) = *γ e*^−*δ τ*^ − *e*^−*δ γ τ*^ .

At the moment of the switch (*τ* = 0), *E*(0) = (*γ* − 1), and the difference of moving averages, *x*(*τ*) − *y*(*τ*), simplifies to equation (8), predicting the sign of the initial rate, *β*_1_. At long times (*τ* → ∞), *E*(*τ*) ≈ 0, and the difference of moving averages again resembles equation (8), but now, instead of predicting *β*_1_, it correctly predicts the sign of the new rate, *β*_2_. In this case, the system exhibits a period of incorrect predictions only when the rate of change of the input signal changes direction. This occurs because the system’s internal model, represented by the two moving averages, requires a finite time to adjust to the new trend.

This adjustment period can be quantified by finding the recovery time *τ*_r_, at which the prediction of the trend direction stops being wrong after the change in the input’s rate of change. Determining the precise recovery time *τ*_r_, where the prediction corrects itself, involves solving a transcendental equation *x*(*τ*) − *y*(*τ*) = 0, but some insight can be gained by focusing on the dynamics in limiting scenarios, or solving the equation numerically (see Figure 2). A small initial rate (|*β*_1_| → 0) leads to a small difference of moving averages, allowing rapid adaptation and a short recovery time (*τ*_r_ → 0). In contrast, a small final rate (|*β*_2_| → 0) results in a slow convergence of the moving averages before the sign of the difference reverses. Consequently, the time required to adapt to the new trend becomes increasingly large, which causes the recovery time to diverge (*τ*_r_ → ∞).

**Figure 2.**
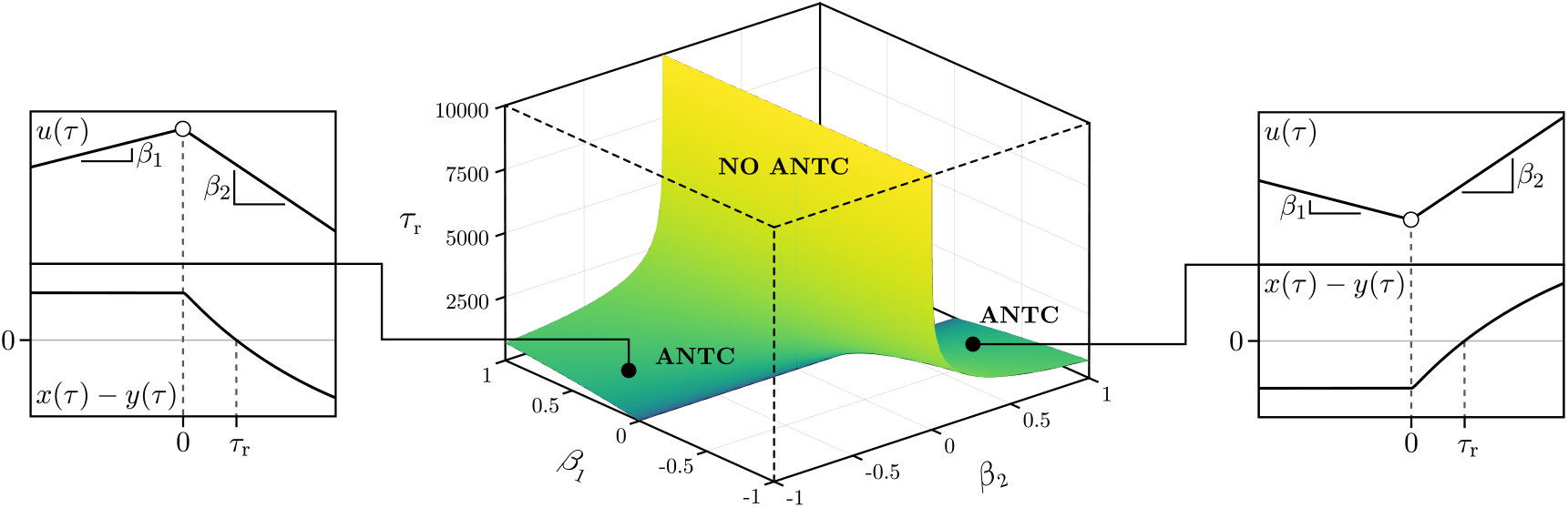
Recovery time. Time *τ*_r_ denotes the recovery time after which trend predictions recover the correct trend after the input’s rate of change abruptly shifts from *β*_1_ to *β*_2_. When the initial and final rates have the same sign (sgn(*β*_1_) = sgn(*β*_2_)), no prediction failures occur; hence *τ*_r_ is undefined, corresponding to the empty regions in the parameter space. When the trend reverses direction, such that *β*_1_ *β*_2_ < 0, a prediction error arises and the recovery time *τ*_r_ is well-defined. The parameter space is symmetric under sign reversal, with the left and right panels illustrating representative examples of the two possible cases: a shift from positive to negative trend (left) and negative to positive trend (right). Parameters were fixed at *α* = 10^−1^, *γ* = 10^−2^, and *δ* = 10^−1^. ANTC: anticipation.

#### 2. Oscillating Input

Our second class of deterministic models considers an important phenomenon, namely the presence of periodic driving inputs associated with environmental cycles (such as daily fluctuations). Let us consider a model of an input signal following

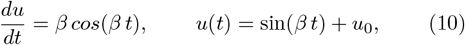

an oscillatory signal with frequency *β* ≥ 0 and initial condition *u*_0_ ≥ 1 to preserve the positivity of concentration.

The asymptotic dynamics of the system as *t* → ∞ are described by the periodic steady-state (see SM I D for full derivation):

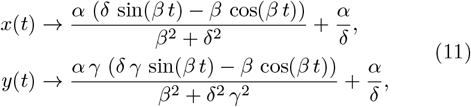

In this case, the difference of moving averages can be expressed as a difference of trigonometric functions

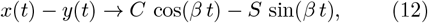

with constant amplitudes defined as

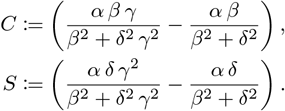

Thus, the predicted trend periodically changes at

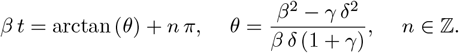

which consists of a phase shift with respect to the cosine sign change from equation (10), with prediction sign in the intervals (*t*_*n*_, *t*_*n*+1_ positive for even *n* and negative for odd *n*.

The union of time spans *T* where predictions are correct, sgn(cos(*β t*)) = sgn(*x*(*t*) − *y*(*t*)), is given by

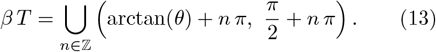

This shows that the system predicts incorrectly immediately after each input trend reversal, exhibiting a delay before recovering accuracy, which increases in magnitude as *β* increases (see Figure 3).

**Figure 3.**
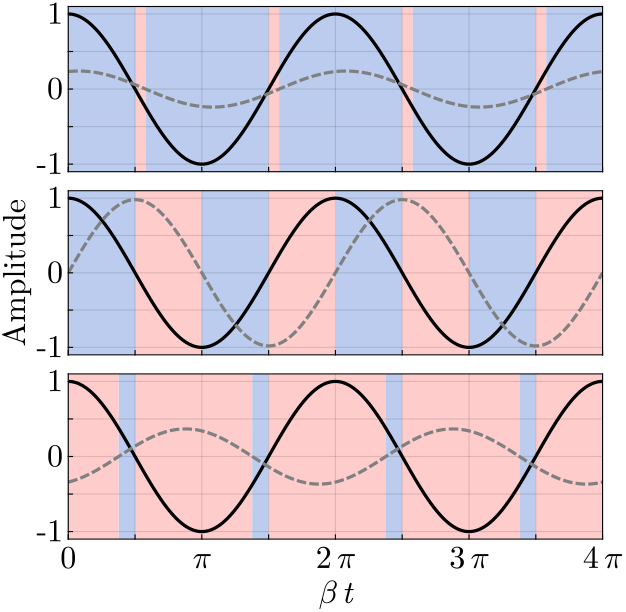
Predicting trends of periodic signals. In black lines, the normalized derivative of the input *u*. In dashed gray lines, the difference of moving averages *x* − *y*. The colored area is blue if the predicted trend is equal to the derivative sign and red otherwise. From top to bottom: *β* = 2.5 · 10^−4^, *β* = 10^−2^, and *β* = 2.5 · 10^−1^. Parameters were fixed at *α* = 10^−1^, *γ* = 10^−2^, and *δ* = 10^−1^.

When the input oscillates slowly, *β* → 0, the input trend and the prediction are in phase. When the input oscillates rapidly, *β* → ∞, the input trend and the prediction are in anti-phase.

### B. Stochastic Input

Understanding the behavior of random processes is crucial in many real-world scenarios, as most signals are not deterministic like those assumed in previous examples. A fundamental starting point for modeling such stochasticity is the Wiener process (also known as Brownian motion), which describes the path of a particle undergoing random movements.

For a standard Wiener process *W* (*t*) with normally distributed, independent increments *dW* (*t*), the conditional probabilities satisfy:

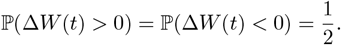

Thus, making any attempt to predict the future direction is no better than a random guess.

Assuming external additive noise for input induction, the system of equations (1) becomes a pair of Ornstein– Uhlenbeck (OU) processes as described by a stochastic differential equation system (SDE),

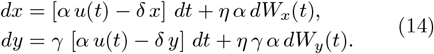

These processes exhibit mean-reverting behavior, with long-term expectations matching the deterministic solutions (*η* = 0),

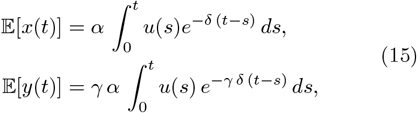

as the expectation of a well-defined Itô integral of an adapted stochastic process is always zero. Meanwhile, the stationary variances remain constant due to noise additivity,

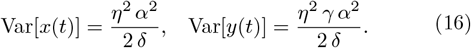

The expected difference of moving averages is straight-forward:

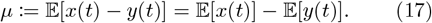

However, computing the variance is more subtle, as it depends on the correlation between the stochastic processes. Assuming that both variables are co-expressed, *dW* (*t*) := *dW*_*x*_(*t*) = *dW*_*y*_(*t*), the stationary variance will converge to

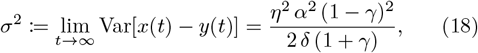

which, as expected, vanishes as *γ* → 1.

Since *u*(*t*) is inherently discrete (e.g., molecule counts in a cell), its granularity becomes significant at low levels. The Wiener process effectively models the accumulation of many small random fluctuations that arise from this discreteness. When *u*(*t*) ≫ 0, the deterministic part of the dynamics dominates, rendering the noise negligible.

In the case of a constant input *u*(*t*) = *u*_0_, the deterministic part of the dynamics vanishes, reducing the predictions to random guessing (*x* − *y*) ∼ 𝒩 (0, *σ*), effectively trying to predict the noise trend *dW* .

#### 1. Constant Drift with Additive Noise

When the system is instead driven by an input with constant drift, like equation (6) in a noisy environment, the expected difference of moving averages *µ* converges to the deterministic solution (8), while stationary the variance *σ*^2^ remains the same as the general case (18) as it is independent of the input (see SM II E.

Although environmental noise introduces variability, the long-term expected prediction remains unbiased and matches the deterministic case. However, individual predictions fluctuate around this expected trend due to stochastic effects.

The probability of a correct trend prediction at any given time is given by

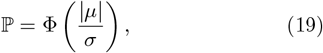

where ϕ(·) is the standard normal cumulative distribution function. As shown in Figure 4a, this probability approaches 1 as |*β*| → ∞ reflecting the dominant position of the drift in the stochastic process, but instead converges to a random guess *P* = 1*/*2 as |*β*| → 0.

**Figure 4.**
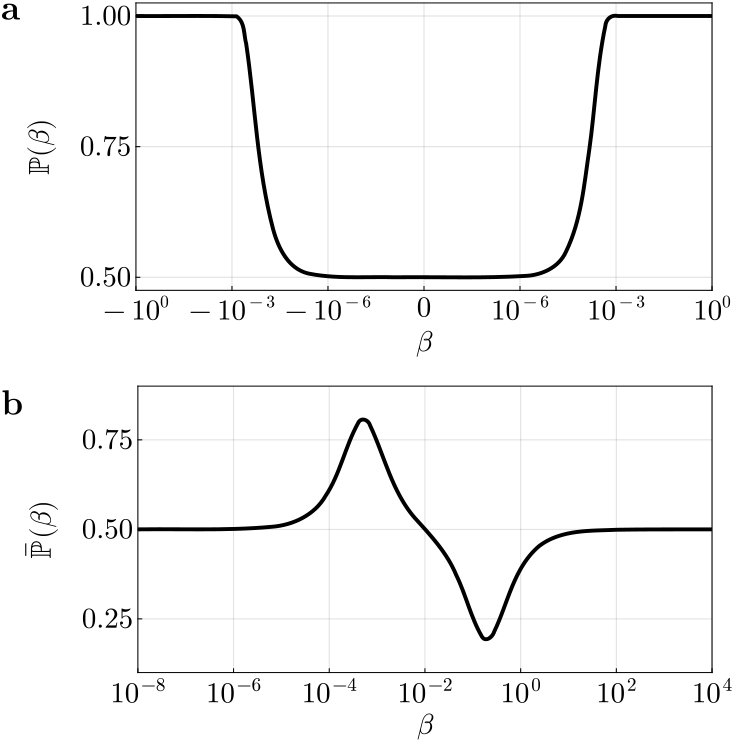
Long-term probabilities of correct prediction under a noisy environment. In (a), the probability of correct trend prediction for an input with a constant drift (19). The *x*-axis is symlog-scaled with a linear region for |*β*| < 10^−8^ to show values outside R^+^ [42]. In (b), the average probability of correct trend prediction over a full period for an oscillatory input (20). Parameters were fixed at *α* = 10^−1^, *γ* = 10^−2^, and *δ* = 10^−1^.

#### 2. Oscillating Input with Additive Noise

Consider the case where the input signal is an oscillating signal (10) in a noisy environment.

Again, the long-term expected value *µ* is given by the deterministic solution (12) and the stationary variance *σ*^2^ given by (18), as noise remains independent of the input (see SM II F.

The probability of correct trend prediction becomes time-dependent,

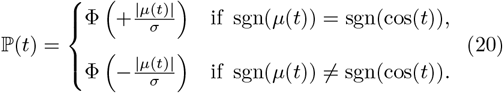

which varies periodically with the input’s derivative.

As illustrated in Figure 4b, the prediction accuracy decays to chance level in both limiting regimes: *β* → 0^+^ and *β* → ∞. This behavior sharply contrasts with the deterministic case, where the system exhibits: (i) asymptotically perfect prediction accuracy as *β* → 0^+^; and (ii) systematically incorrect predictions as *β* → ∞. The system enters a noise-governed phase in both limits, as the deterministic contribution vanishes, effectively randomizing the output.

#### 3. Exponentially Weighted Moving Averages

Consider a Wiener process *u* = {*W* (*t*_*k*_)} sampled at equally spaced time points,

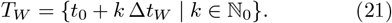

with a time step of size Δ*t*_*W*_ . To introduce predictability, a transformed process *û* is constructed via exponentially weighted moving averages (EWMA) [43]. The transformation applies time-series smoothing by assigning exponentially decreasing weights to past observations, emphasizing recent data, while retaining the diminishing influence of past values.

The transformed process at the discrete time points *t*_*k*_ ∈ *T*_*W*_ is computed using EWMA as follows:

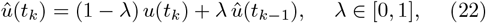

with initial condition *û*(*t*_0_) = *u*(*t*_0_).

A continuous-time approximation *û*(*t*) for *t* ∉ *T*_*W*_ is then obtained by linear interpolation,

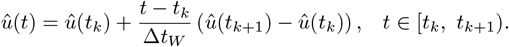

Here, the input signal is precomputed and shifted so that no negative concentrations are present.

To numerically evaluate the system, stochastic simulations are conducted with zero initial conditions and allowed to evolve until sufficient time has elapsed for transient dynamics to become negligible (see Figure 5a as an example). The true trend, denoted as

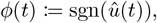

and the predicted trend, namely

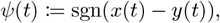

can then be computed and compared to assess system performance.

**Figure 5.**
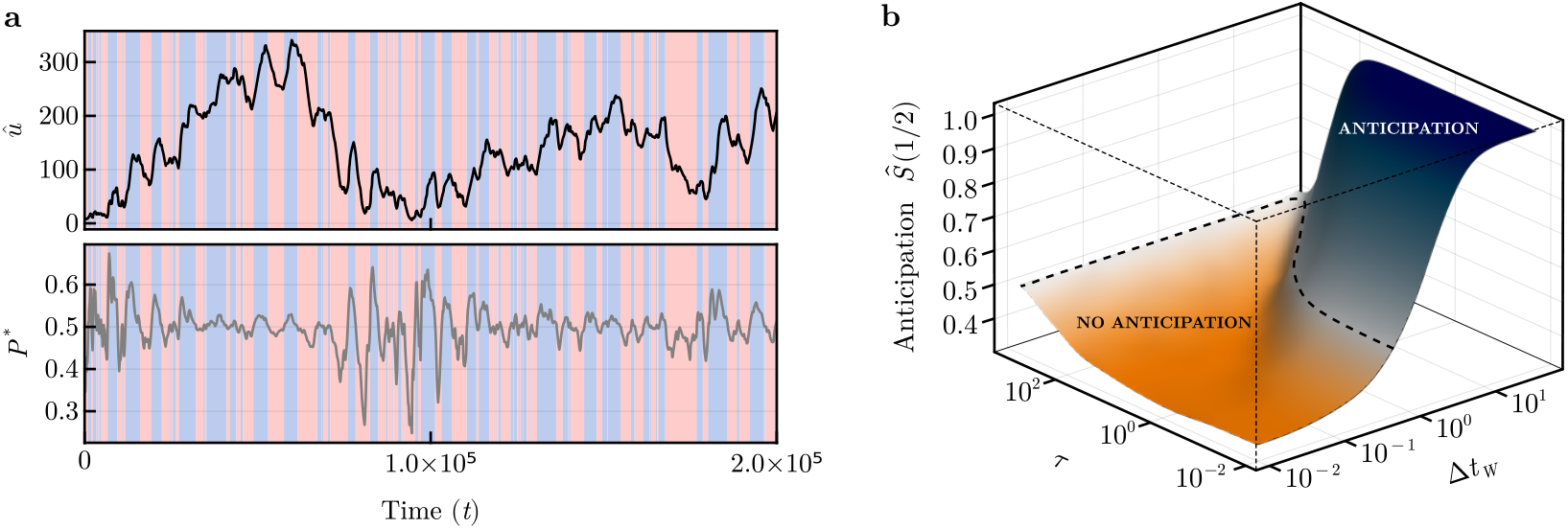
Trend prediction of EWMA-transformed Wiener processes. In (a), a sample time-series from a simulation. In black, the input signal *û*, and in gray the output observable *P* ^∗^. The background is colored depending on the level of *P* ^∗^: blue if the trend is predicted to be positive (*x* − *y* > 0), red if the prediction is negative (*x* − *y* < 0). In (b), a surface showing the expected fraction of time-series where the predictions are expected to be at least as good as a random guesser, *Ŝ*_*M*_ = 1*/*2. The dashed line at the 1*/*2 level represents the expected outcome of a random guesser (see SM II B, II C). The divergent color map is centered around this value to show how much the expectation differs. Parameters were fixed at *α* = 10^−1^, *γ* = 10^−2^, *δ* = 10^−1^, *λ* = 2 · 10^−1^, and *M* = 10^6^.

Of particular interest are predictions of the trend at time *t* for a future time *t* + *τ*, as any downstream effector requires a finite time interval to exert its influence.

The prediction accuracy *A*(*τ*) quantifies the empirical probability that the system’s predicted trend at a given time *ψ*(*t*_*i*_) correctly anticipates the true trend *ϕ*(*t*_*i*_ +*τ*) at a time horizon *τ* . It can be computed over *N* uniformly sampled time points 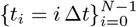 as

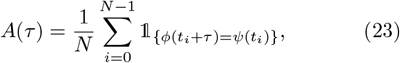

where 𝟙 _{·}_ is an indicator function which returns 1 if the condition is true and 0 otherwise. This measure computes a normalized, shifted cross-correlation over a time-series, with *A*(*τ*) = 1 indicating perfect prediction, *A*(*τ*) = 1*/*2 corresponding to random guessing, and *A*(*τ*) = 0 signifying complete opposition to the true trend.

To assess system robustness, revealing whether prediction accuracy is consistently achievable or heavily dependent on specific realizations of the input, an ensemble of *M* simulations with different stochastic trajectories for the input can be performed. The empirical cumulative distribution function (eCDF) of the prediction accuracy across the ensemble can be estimated following Vaart [44] as

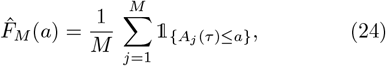

where parameter *a* represents a threshold accuracy level of interest, allowing one to compute the fraction of simulations in the ensemble that achieve an accuracy of at most *a*. To assess the probability that the system will exceed the threshold, the empirical survival function can be derived as the complement of the eCDF:

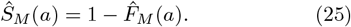

This function provides a direct measure of robustness: If *Ŝ*_*M*_ (1*/*2) > 1*/*2, an individual is expected to perform better than random, while *Ŝ*_*M*_ (1*/*2) < 1*/*2 indicates that individuals are expected to perform systematically worse than random.

Moving beyond purely independent predictions, an alternative system could be developed where individuals exchange information and work towards a consensus before making a collective prediction, a concept inspired by biological *quorum sensing* [45, 46]. This collective approach can potentially lead to more robust decisions in some cases [47, 48], but would inevitably introduce some additional delay.

In such a system, each cell would still process input and produce an individual prediction, represented as a diffusible molecule detectable by other cells. A collective prediction would only be generated if a *quorum*—a minimum concentration of this molecule in the environment— is reached.

Under the assumption of independent predictions by individual cells, the proposed system mirrors the core concepts of an extended Condorcet jury theorem, where heterogeneous voters with different competence levels make collective decisions [49, 50]. Within this framework, each simulation in an ensemble can be viewed as an independent “voter” and the prediction accuracy corresponds to the voter’s competence.

In the cases where *Ŝ*_*M*_ (1*/*2) > 1*/*2, the ensemble’s average probability of correct prediction is also better than random chance, mirroring the scenario where voters are competent on average. Statistical consensus naturally enhances robustness, as majority voting improves accuracy compared with individual predictions. In contrast, if the average individual prediction is worse than random, aggregated predictions underperform both random guessing and individual estimates.

These two regimes are illustrated in Figure 5b. At short input autocorrelation times (Δ*t*_*W*_ → 0), the system struggles to adapt to rapid fluctuations, resulting in expected performance worse than random. As autocorrelation increases, the system achieves a higher expected prediction accuracy and maintains reliable predictions into the future. However, since the input correlations remain local in time, accuracy decays to random levels as the prediction horizon increases (*τ* → ∞).

## IV. DISCUSSION

Anticipation is considered a crucial component of cognitive complexity. In general terms, it involves the use of past and present information to make possible predictions about future events. The concept has traditionally been tied to human psychology and the behavior of neural agents, but this view has changed, embracing organisms and systems of many different kinds. Anticipation can be found across scales in biology [9], including aneural organisms [10]. An important challenge in all these cases is to properly define a measure of anticipation that can help avoid ambiguities regarding the nature and efficiency of the prediction, particularly in stochastic environments.

In this paper, we have taken a different path to explore the problem of anticipation by considering how to build a minimal circuit that can implement trend prediction within a cell. The starting point is Frank’s model [27, 51], which we explicitly map to a specific genetic motif. The goal here is twofold: to provide formally well-defined, engineerable designs that can be practically implemented, and to quantitatively assess the conditions under which anticipatory behavior is expected to emerge. This has been done both mathematically and numerically, showing that anticipation is an expected property of this system. In all the considered cases, there is a well-defined domain where anticipation is shown to be present, and the bounds are precisely determined.

Given the simplicity of our circuit design and its potential alternatives, we conjecture that anticipatory behavior (with different levels of accuracy) might be widespread in cellular networks. Indeed, natural systems already employ analogous predictive strategies. For instance, bacterial chemotaxis uses temporal sampling and precise adaptation to approximate spatial gradients without spatial sensing [52, 53]. Similarly, bacterial communities could exploit *quorum sensing* compounds with different diffusion rates to increase the information yield about their environment, effectively generating multi-timescale signaling analogous to that arising from differential degradation in the proposed circuits [54–56] . More explicit implementations also exist, such as two-component systems leveraging production coupled to environmental dynamics and differential decay [57]. When cellular response timescales are mismatched to environmental signal dynamics or sensing is overwhelmed by noise, cells may adopt probabilistic strategies such as “bet-hedging”, bypassing the need for precise sensing [58–61].

Beyond natural systems, the potential for the design of novel circuits to be engineered within living cells opens the door to studying the evolution of synthetic behavior and its evolutionary dynamics [62]. Further extensions of this work should consider the coupling between single- and multicellular scales. In this context, both microbial communities [18, 63] and tissues [12, 64, 65] display collective intelligence characteristics that could be enhanced through synthetic biology [41].

## ACKNOWLEDGMENTS

The authors thank the members of the Complex Systems Lab for their valuable discussions and insights and the Santa Fe Institute. They also thank Carlos Greykey for inspiring ideas.

This work has been supported by the AGAUR 2021 SGR 0075 grant and the Santa Fe Institute.

J. P. M. was supported by grant PRE2020-091968, funded by MCIN/AEI (10.13039/501100011033), and co-funded by the ESF through the program “Investing in your future”.

