## Supplementary Material for "A Minimal Genetic Circuit for Cellular Anticipation"

### I. DETERMINISTIC DYNAMICS

Given an input signal $u$, the moving averages $x$ and $y$, can be modeled as

$$\begin{aligned}\frac{dx}{dt} &= \alpha\, u(t) - \delta\, x,\\ \frac{dy}{dt} &= \gamma\left(\alpha\, u(t) - \delta\, y\right),\end{aligned} \tag{1}$$

where $\alpha \geq 0$ is a production rate in response to the input signal, $\delta \geq 0$ represents the degradation/dilution rate, and $0 < \gamma < 1$ is a scaling factor that makes one gene respond more slowly than the other, creating the different timescales.

The equation for $y(t)$ is a linear first-order ODE of the form

$$\frac{dy}{dt} + P(t)\, y = Q(t),$$

where $P(t) = \gamma\,\delta$ and $Q(t) = \gamma\,\alpha\, u(t)$, with an integrating factor

$$M(x) = e^{\int P(t)\, dt} = e^{\int \gamma\,\delta\, dt} = e^{\gamma\,\delta\, t}.$$

To solve the differential equation, start by multiplying both sides of the ODE by the integrating factor:

$$e^{\gamma\,\delta\, t}\frac{dy}{dt} + \gamma\,\delta\, e^{\gamma\,\delta\, t} y = \gamma\,\alpha\, u(t)\, e^{\gamma\,\delta\, t},$$

which makes the left hand side the derivative of $y(t)\, e^{\gamma\,\delta\, t}$:

$$\frac{d}{dt}\left(y(t)\, e^{\gamma\,\delta\, t}\right) = \gamma\,\alpha\, u(t)\, e^{\gamma\,\delta\, t}.$$

Integrate both sides with respect to $t$:

$$y(t)\, e^{\gamma\,\delta\, t} = \gamma\,\alpha \int_0^t u(s)\, e^{\gamma\,\delta\, s}\, ds + y_0,$$

where $y_0$ is the constant of integration (initial condition).

Solve for $y(t)$:

$$y(t) = e^{-\gamma\,\delta\, t}\left(\gamma\,\alpha \int_0^t u(s)\, e^{\gamma\,\delta\, s}\, ds + y_0\right). \tag{2}$$

The solution for $x(t)$ is the same as for $y(t)$ when $\gamma = 1$. Therefore, $\gamma = 1$ can be directly substituted into equation (2):

$$x(t) = e^{-\,\delta\, t}\left(\alpha \int_0^t u(s)\, e^{\delta\, s}\, ds + x_0\right). \tag{3}$$

#### A. Heaviside Input

The simplest dynamical input is an a piecewise constant function, a fundamental building block for representing and analyzing signals [1]:

$$u(t) = \begin{cases} u_1, & t < 0, \\ u_2, & t \geq 0. \end{cases} \tag{4}$$

For $t < 0$, the system is in steady state with $u(t) \coloneqq u_1$. The steady-state solutions are

$$\frac{dx}{dt} = 0 \implies x = \frac{\alpha\, u_1}{\delta},$$
$$\frac{dy}{dt} = 0 \implies y = \frac{\alpha\, u_1}{\delta}.$$

Thus, at $t = 0$, the initial conditions are known:

$$x(0) = \frac{\alpha\, u_1}{\delta}, \qquad y(0) = \frac{\alpha\, u_1}{\delta}.$$

For $t \geq 0$,

$$y(t) = e^{-\gamma\,\delta\,t}\left(\gamma\,\alpha\,u_2 \int_0^t e^{\gamma\,\delta\,s} ds + y(0)\right),$$

which can be evaluated by substituting the initial condition and the integral

$$\int_0^t e^{\gamma\,\delta\,s}\, ds = \frac{e^{\gamma\,\delta\,t} - 1}{\gamma\,\delta}.$$

Thus, the solution becomes

$$y(t) = \frac{\alpha\, u_2}{\delta}(1 - e^{-\gamma\,\delta\,t}) + \frac{\alpha\, u_1}{\delta} e^{-\gamma\,\delta\,t}.$$

And $x(t)$ can be easily computed by substituting $\gamma = 1$:

$$x(t) = \frac{\alpha\, u_2}{\delta}(1 - e^{-\delta\,t}) + \frac{\alpha\, u_1}{\delta} e^{-\delta\,t},$$

which gives an expression for the difference of moving averages as

$$x(t) - y(t) = \frac{\alpha\, u_2}{\delta}\left(e^{-\gamma\,\delta\,t} - e^{-\delta\,t}\right) + \frac{\alpha\, u_1}{\delta}\left(e^{-\delta\,t} - e^{-\gamma\,\delta\,t}\right).$$

The system maintains equilibrium $(x - y) = 0$ both before the input change ($t < 0$) and after full adaptation ($t \to \infty$). During the transient phase following an input discontinuity at $t = 0$, $\text{sgn}(x(t) - y(t))$ provides immediate directional information about the change:

- If the input suddenly increases ($u_2 > u_1$), the trend is predicted as positive ($x - y > 0$).
- If the input suddenly decreases ($u_2 < u_1$), the trend is predicted as negative ($x - y < 0$).

Remarkably, this trend detection occurs despite the input's discontinuous jump, demonstrating how the system's dual-timescale dynamics encodes directional information without requiring a well-defined derivative of the input signal (see an example in Figure 1).

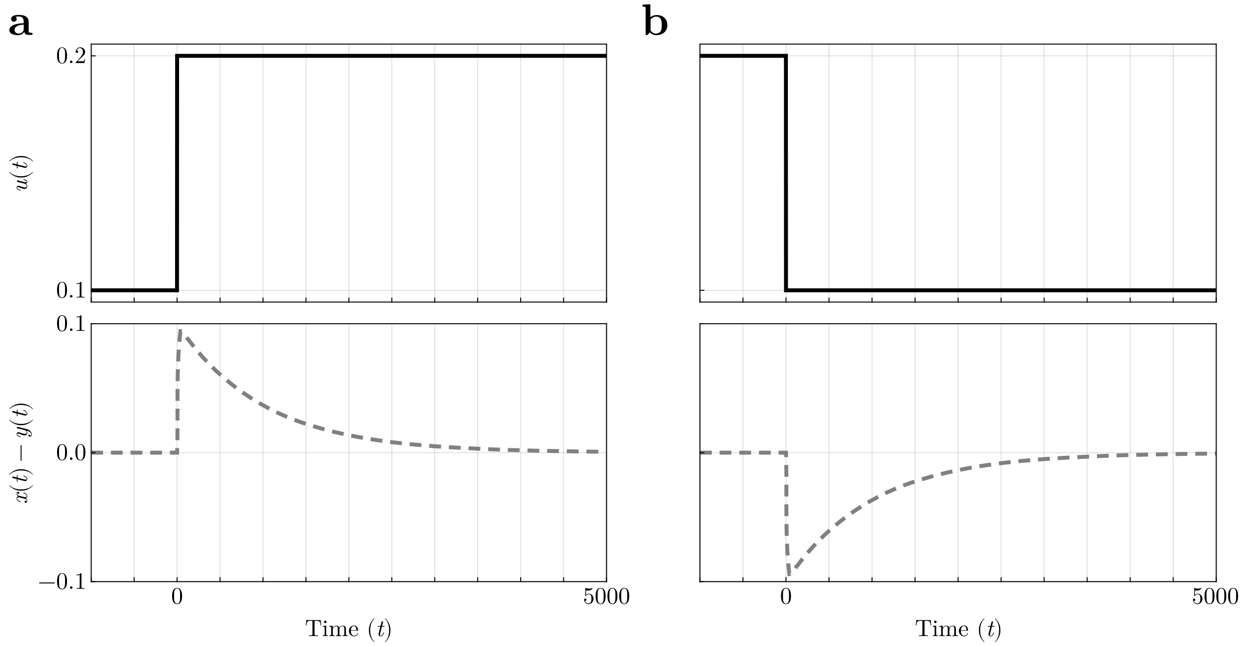


Figure 1. Sample time-series of the system under a piecewise constant input. In (a), the behavior when $u_1 < u_2$, where the system predicts a trend decrease during the transient dynamics. In (b), the behavior when $u_1 > u_2$, where the system predicts a trend increase during the transient dynamics. In both cases, when $t \to \infty$ the system returns to the steady state for a constant input, $(x - y) = 0$. Parameters were fixed at $\alpha = 10^{-1}$, $\gamma = 10^{-2}$, and $\delta = 10^{-1}$.

#### B. Constant Drift Input

Given an input with a constant rate of change:

$$\frac{du}{dt} = \beta, \qquad u(t) = \beta\, t + u_0, \tag{5}$$

the solution of the system of equations (1) is given by

$$\begin{aligned} x(t) &= \frac{\alpha\,(\beta\, t + u_0)}{\delta} - \frac{\alpha\,\beta}{\delta^2} + \left(\frac{\alpha\,\beta}{\delta^2} - \frac{\alpha\, u_0}{\delta} + x_0\right) e^{-\delta\, t}, \\ y(t) &= \frac{\alpha\,(\beta t + u_0)}{\delta} - \frac{\alpha\,\beta}{\delta^2\,\gamma} + \left(\frac{\alpha\,\beta}{\delta^2\,\gamma} - \frac{\alpha\, u_0}{\delta} + y_0\right) e^{-\delta\,\gamma\, t}. \end{aligned} \tag{6}$$

Thus, the trend will be anticipated as positive when $(x - y) > 0$, and negative otherwise:

$$x(t) - y(t) = \frac{\alpha\,\beta}{\delta^2\,\gamma}\,(1-\gamma) + \left(\frac{\alpha\,\beta}{\delta^2} - \frac{\alpha\, u_0}{\delta} + x_0\right) e^{-\delta\, t} - \left(\frac{\alpha\,\beta}{\delta^2\,\gamma} - \frac{\alpha\, u_0}{\delta} + y_0\right) e^{-\delta\,\gamma\, t}. \tag{7}$$

At long times ($t \to \infty$), the expression has the same sign as the slope of the input:

$$\lim_{t\to\infty} x(t) - y(t) = \frac{\alpha\,\beta}{\delta^2\,\gamma}\,(1-\gamma)\,. \tag{8}$$

#### C. Piecewise Constant Drift Input

An extension of this dynamics involves an input that changes at a constant rate $\beta_1$ in $t \in [t_0, t_s)$, and at some point starts changing at a rate $\beta_2$ for $t \geq t_s$. This is given by a piecewise input function:

$$u(t) = \begin{cases} \beta_1\, t + u_0, & t < t_s, \\ \beta_2\,(t - t_s) + \beta_1\, t_s + u_0, & t \geq t_s. \end{cases}$$

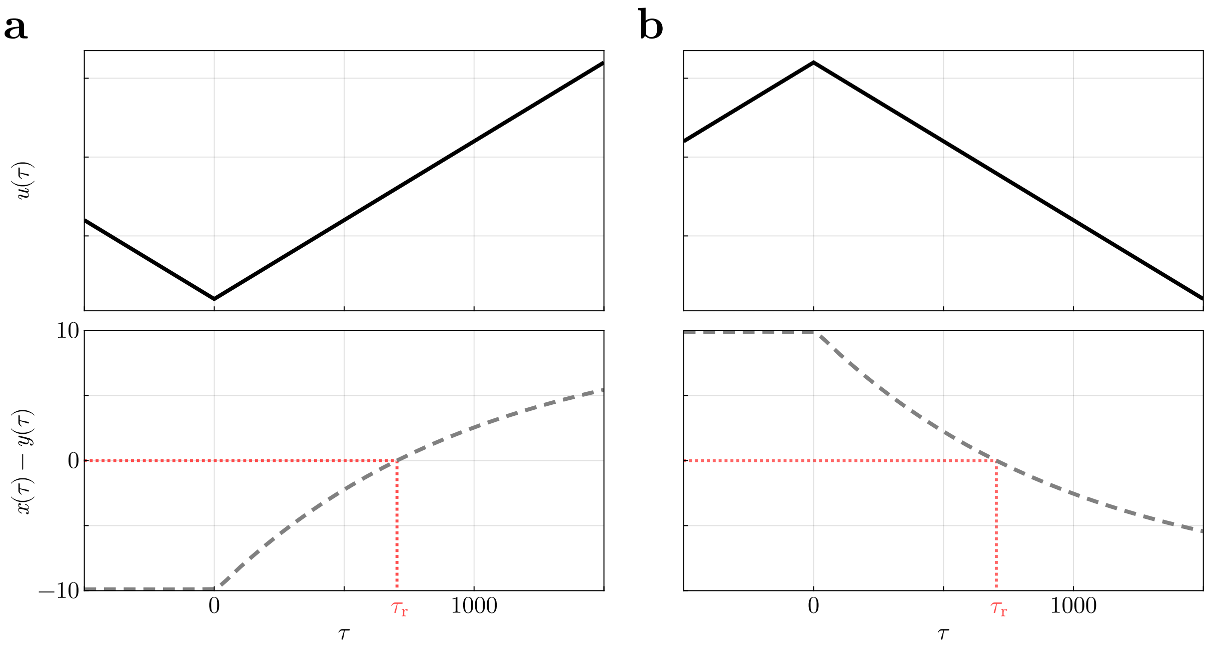


Figure 2. Sample time-series of the system under an input with constant drift, where the drift direction reverses at $\tau = 0$. In (a), where $\beta_1 < 0$ and $\beta_2 > 0$, the system initially retains the sign of $\beta_1$, yielding $(x - y) < 0$ for a transient period $(0,\ \tau_{\mathrm{r}})$. Beyond this interval, the difference in means grows monotonically, approaching the steady state predicted by expression (eq:piecewise-drift-longterm-difference). In (b), where $\beta_1 > 0$ and $\beta_2 < 0$, the dynamics follow a symmetric pattern: the system first predicts an upward trend before transitioning to a decline, mirroring the behavior in (a) but with opposing drift directions. Parameters were fixed at $\alpha = 10^{-1}$, $\gamma = 10^{-2}$, and $\delta = 10^{-1}$.

The solution of the system for $t < t_s$ is the same as system (6), and for $t \geq t_s$,

$$\begin{aligned} x(t) &= \frac{\alpha\ (\beta_2\, t + (\beta_1 - \beta_2)\, t_s + u_0)}{\delta} - \frac{\alpha\, \beta_2}{\delta^2} + \frac{\alpha\, (\beta_2 - \beta_1)}{\delta^2}\, e^{-\delta\, (t - t_s)} + \left( \frac{\alpha\, \beta_1}{\delta^2} - \frac{\alpha\, u_0}{\delta} + x_0 \right) e^{-\delta\, t}, \\ y(t) &= \frac{\alpha\ (\beta_2 t + (\beta_1 - \beta_2)\, t_s + u_0)}{\delta} - \frac{\alpha\, \beta_2}{\delta^2\, \gamma} + \frac{\alpha\, (\beta_2 - \beta_1)}{\delta^2\, \gamma}\, e^{-\delta\, \gamma\, (t - t_s)} + \left( \frac{\alpha\, \beta_1}{\delta^2\, \gamma} - \frac{\alpha\, u_0}{\delta} + y_0 \right) e^{-\delta\, \gamma\, t}. \end{aligned} \tag{9}$$

Assuming the switch takes place at time $t_s$ sufficiently large such that the transient dynamics before the event are negligible, and taking $\tau \coloneqq t - t_s$,

$$\begin{aligned} x(\tau) &\approx \frac{\alpha\ (\beta_2\, (\tau + t_s)) + (\beta_1 - \beta_2)\, t_s + u_0)}{\delta} - \frac{\alpha\, \beta_2}{\delta^2} + \frac{\alpha\, (\beta_2 - \beta_1)}{\delta^2}\, e^{-\delta\, \tau}, \\ y(\tau) &\approx \frac{\alpha\ (\beta_2 (\tau + t_s) + (\beta_1 - \beta_2)\, t_s + u_0)}{\delta} - \frac{\alpha\, \beta_2}{\delta^2\, \gamma} + \frac{\alpha\, (\beta_2 - \beta_1)}{\delta^2\, \gamma}\, e^{-\delta\, \gamma\, \tau}. \end{aligned} \tag{10}$$

Thus, the difference of moving averages becomes

$$x(\tau) - y(\tau) \approx \frac{\alpha}{\delta^2\, \gamma}\ \left( \beta_2\, (1 - \gamma) + (\beta_2 - \beta_1)\ \left( \gamma\, e^{-\delta\, \tau} - e^{-\delta\, \gamma\, \tau} \right) \right). \tag{11}$$

As observed, the system undergoes a phase of incorrect predictions when the input signal's rate of change shifts direction, requiring a finite adjustment period to align with the new trend.

This adjustment duration can be quantified by determining the recovery time $\tau_{\mathrm{r}}$, which marks the point after the input's rate change when the system's trend predictions are no longer erroneous.

The recovery time $\tau_{\mathrm{r}}$ can be obtained by solving the transcendental equation

$$\frac{\alpha}{\delta^2\, \gamma}\ \left( \beta_2\, (1 - \gamma) + (\beta_2 - \beta_1)\ \left( \gamma\, e^{-\delta\, \tau} - e^{-\delta\, \gamma\, \tau} \right) \right) = 0. \tag{12}$$

Although this expression typically has no closed-form solution, numerical methods provide a practical approach (see Figure 2). However, analytical conditions for the existence of a zero can still be derived. Furthermore, for minor perturbations, the recovery time can be estimated via a Taylor expansion of the equation. These topics are examined in detail in the subsequent sections.

*Existance and Uniqueness of a Recovery Time*

The conditions under which a zero will exist for equation (12) given a pair of rates ($\beta_1,\ \beta_2$) can be analytically derived as follows.

Define a function $f(\tau)$, which determines the sign of the difference of moving averages from equation (11):

$$f(\tau) \coloneqq \beta_2\,(1-\gamma) + (\beta_2 - \beta_1)\,\left(\gamma\, e^{-\delta\,\tau} - e^{-\delta\,\gamma\,\tau}\right), \qquad \tau \geq 0. \tag{13}$$

The behavior of $f(\tau)$ exhibits the following behavior at $\tau = 0$ and as $\tau \to \infty$:

- At $\tau = 0$, the expression is the steady state prediction of a constant rate input $\beta_1$:

$$f(0) = \beta_1\,(1-\gamma).$$

- As $\tau \to \infty$, the expression is the steady state prediction of a constant rate input $\beta_2$:

$$\lim_{\tau\to\infty} f(\tau) = \beta_2\,(1-\gamma),$$

Since $(1-\gamma) > 0$ the signs of the expressions are determined by the sign of $\beta_1$ and $\beta_2$ respectively.

If $\mathrm{sgn}(\beta_1) \neq \mathrm{sgn}(\beta_2)$, then $f(0)$ and $\lim_{\tau\to\infty} f(\tau)$ also have opposite signs. Since $f(\tau)$ is continuous, by the Intermediate Value Theorem, there must be at least one zero in the interval $\tau \in [0, \infty)$.

If $\mathrm{sgn}(\beta_1) = \mathrm{sgn}(\beta_2)$, then $f(0)$ and $\lim_{\tau\to\infty} f(\tau)$ have the same sign. In this case, for a zero to exist, there must be an even number of zeros in the interval $\tau \in [0, \infty)$.

To demonstrate that if there is a zero it must be unique, monotonicity of the function $f(\tau)$ can be analyzed by computing its derivative with respect to $\tau$ as

$$\frac{df(\tau)}{d\tau} = \delta\,\gamma\,(\beta_2 - \beta_1)\,\left(e^{-\delta\,\gamma\,\tau} - e^{-\delta\,\tau}\right).$$

Since $\gamma \in (0,\ 1) \implies \left(e^{-\delta\,\gamma\,\tau} - e^{-\delta\,\tau}\right) > 0$, the sign of the derivative is fully determined by the sign of $(\beta_2 - \beta_1)$, being strictly increasing if $\beta_2 > \beta_1$ and strictly decreasing if $\beta_2 < \beta_1$. In either case, $f(\tau)$ is monotonic. Therefore, if there is a zero, it must be unique.

Combining the existence and uniqueness results, it can be seen that $f(\tau)$ has exactly one zero if and only if $\mathrm{sgn}(\beta_1) \neq \mathrm{sgn}(\beta_2)$. If $\mathrm{sgn}(\beta_1) = \mathrm{sgn}(\beta_2)$, then $f(\tau)$ has no zeros. This means that a sign change in the input rate is a necessary and sufficient condition for the existence of a time interval where the predicted trend is reversed.

*Taylor Series Approximation*

For small values of $\tau$, the value of $e^{a\,\tau}$ can be approximated by keeping only the first terms of the Taylor series:

$$e^{a\,\tau} = \sum_{n=0}^{\infty} \frac{f^{(n)}(0)}{n!}\,\tau^n = \sum_{n=0}^{\infty} \frac{a^n}{n!}\,\tau^n = 1 + a\,\tau + \frac{a^2\,\tau^2}{2!} + \frac{a^3\,\tau^3}{3!} + \ldots,$$

thus, with a second order approximation:

$$\gamma\, e^{-\delta\,\tau} - e^{-\delta\,\gamma\,\tau} \approx \gamma - 1 + \frac{\delta^2\,\gamma\,\tau^2}{2}\,(1-\gamma) + O\left(\tau^3\right).$$

Substituting into the difference of moving averages from equation (11), the difference becomes

$$x(\tau) - y(\tau) \approx \frac{\alpha}{\delta^2\,\gamma}\,\left(\beta_2\,(1-\gamma) + (\beta_2-\beta_1)\,\left(\gamma - 1 + \frac{\delta^2\,\gamma\,\tau^2}{2}\,(1-\gamma)\right)\right),$$

which has two solutions for $\tau$,

$$\tau_{\pm} = \pm\sqrt{\frac{2\,\beta_1 + O\left(\tau^3\right)}{\delta^2\,\gamma\,(\beta_1-\beta_2)}},$$

In this case, the only valid value is $\tau_+ > 0$ due to the piecewise nature of the problem being treated.

While this approximation can be useful in some cases, the computed value can be wildly different than the true value. Thus, it is advisable to use numerical methods to approximate the solution instead.

#### D. Oscillating Input

Given an oscillating input signal

$$\frac{du}{dt} = \beta\,\cos(\beta\,t), \qquad\qquad u(t) = 1 + \sin(\beta\,t), \tag{14}$$

with frequency $\beta > 0$, the solution of the system of equations (1) is given by

$$\begin{aligned} x(t) &= \frac{\alpha\,\left(\delta\,\sin(\beta\,t) - \beta\,\cos(\beta\,t)\right)}{\beta^2+\delta^2} + \frac{\alpha}{\delta} - \left(\frac{\alpha\,\left(\beta^2 - \beta\,\delta + \delta^2\right)}{\delta\,\left(\beta^2+\delta^2\right)} - x_0\right)\,e^{-\delta\,t}, \\ y(t) &= \frac{\alpha\,\gamma\,\left(\delta\,\gamma\,\sin(\beta\,t) - \beta\,\cos(\beta\,t)\right)}{\beta^2+\delta^2\,\gamma^2} + \frac{\alpha}{\delta} - \left(\frac{\alpha\,\left(\beta^2 - \beta\,\delta\,\gamma + \delta^2\,\gamma^2\right)}{\delta\,\left(\beta^2+\delta^2\,\gamma^2\right)} - y_0\right)\,e^{-\delta\,\gamma\,t}. \end{aligned} \tag{15}$$

Given the known derivative of the input from equation (14), the trigonometric function determines the intervals at which the trend is positive or negative:

$$\operatorname{sgn}\left(\beta\,\cos(\beta\,t)\right) = (-1)^n, \qquad \beta\,t \in \left(n\,\pi - \frac{\pi}{2},\; n\,\pi + \frac{\pi}{2}\right), \qquad n \in \mathbb{Z}.$$

After a sufficiently long time has elapsed such that transient effects from initial conditions have decayed, the difference between the moving averages settles into a periodic steady state given by

$$x(t) - y(t) \to \left(\frac{\alpha\,\beta\,\gamma}{\beta^2+\delta^2\,\gamma^2} - \frac{\alpha\,\beta}{\beta^2+\delta^2}\right)\cos(\beta\,t) - \left(\frac{\alpha\,\delta\,\gamma^2}{\beta^2+\delta^2\,\gamma^2} - \frac{\alpha\,\delta}{\beta^2+\delta^2}\right)\sin(\beta\,t). \tag{16}$$

To find the zeros of this expression, the coefficients can be further simplified by using trigonometric functions. Let $C$ be the coefficient of $\cos(\beta\,t)$ and $S$ be the coefficient of $\sin(\beta\,t)$. After factoring and simplifying

$$C := \alpha\,\beta\,\frac{(1-\gamma)(\gamma\,\delta^2 - \beta^2)}{(\beta^2+\delta^2\,\gamma^2)(\beta^2+\delta^2)}, \qquad\qquad S := \alpha\,\delta\,\frac{-\beta^2\,(1-\gamma)(1+\gamma)}{(\beta^2+\delta^2\,\gamma^2)(\beta^2+\delta^2)}.$$

Substituting these into the original equation and simplifying yields the following equality:

$$\frac{\alpha\,\beta\,(1-\gamma)}{(\beta^2+\delta^2\,\gamma^2)(\beta^2+\delta^2)}\,\left[(\gamma\,\delta^2 - \beta^2)\,\cos(\beta\,t) + \beta\,\delta\,(1+\gamma)\,\sin(\beta\,t)\right] = 0,$$

which is a constant nonzero positive factor multiplied by a trigonometric time-dependent expression.

Assuming $\cos(\beta\,t) \neq 0$, as $cos(x) = 0 \implies \sin(x) = \pm 1$, the expression can be rewritten as

$$\tan(\beta\,t) = \frac{\beta^2 - \gamma\,\delta^2}{\beta\,\delta\,(1+\gamma)},$$

which has periodic solutions at

$$\beta\,t = \arctan\left(\frac{\beta^2 - \gamma\,\delta^2}{\beta\,\delta\,(1+\gamma)}\right) + n\,\pi, \qquad n \in \mathbb{Z}.$$

The prediction sign can be then computed by computing the derivative of equation (16),

$$\frac{d}{dt}\,(x(t) - y(t)) \approx \frac{\alpha\,\beta^2\,(1-\gamma)}{(\beta^2 + \delta^2\,\gamma^2)(\beta^2+\delta^2)}\left[(\beta^2 - \gamma\,\delta^2)\,\sin(\beta\,t) + \beta\,\delta\,(1+\gamma)\,\cos(\beta\,t)\right],$$

which has the following form:

$$C\,\left[A\,\sin(\beta\,t) + B\,\cos(\beta\,t)\right],$$

where

$$A := \beta^2 - \gamma\,\delta^2, \qquad B := \beta\,\delta\,(1+\gamma), \qquad C := \frac{\alpha\,\beta^2\,(1-\gamma)}{(\beta^2 + \delta^2\,\gamma^2)(\beta^2+\delta^2)}.$$

The term $A\sin(\beta t) + B\cos(\beta t)$ can be rewritten using a single trigonometric function with a phase shift. Recall that

$$A\,\sin(\beta\,t) + B\,\cos(\beta\,t) = \sqrt{A^2+B^2}\,\sin(\beta\,t + \phi), \qquad \phi = \arctan\left(\frac{B}{A}\right);$$

Geometrically, the coefficients $A$ and $B$ can be interpreted as the two sides of a right triangle. The resulting oscillation amplitude corresponds to the length of the hypotenuse, namely

$$h = \sqrt{A^2+B^2} = \sqrt{(\beta^2 - \gamma\,\delta^2)^2 + (\beta\,\delta\,(1+\gamma))^2}.$$

Thus, the derivative becomes

$$\frac{d}{dt}\,(x(t) - y(t)) \approx C\,h\,\sin(\beta\,t + \phi).$$

The expression $\sin(\beta\,t + \phi)$ oscillates between $-1$ and 1. To find the maximal magnitude of the derivative (which determines the sign of the prediction), it can be evaluated at times $t_n$ where reaches its extrema:

$$\sin(\beta\,t_n + \phi) = (-1)^n, \qquad \beta\,t_n = \frac{\pi}{2} + n\,\pi - \phi, \qquad n \in \mathbb{Z}.$$

At these times, the derivative simplifies to

$$\frac{d}{dt}\,(x(t) - y(t)) \approx C\,h\,(-1)^n.$$

Substituting $C$, the derivative becomes

$$\frac{d}{dt}\,(x(t) - y(t)) \approx (-1)^n\,\frac{\alpha\,\beta^2\,(1-\gamma)}{(\beta^2 + \delta^2\,\gamma^2)(\beta^2+\delta^2)}\,h.$$

with $h$ previously defined as

$$h = \sqrt{\left(\beta^2 - \gamma\,\delta^2\right)^2 + \left(\beta\,\delta\,(1+\gamma)\right)^2}.$$

As all terms are positive except $(-1)^n$, an interval $(t_n,\ t_{n+1})$ is positive for even $n$ and negative for odd $n$.

The union of time spans $T$ where predictions are correct, $\mathrm{sgn}(\cos(\beta\,t)) = \mathrm{sgn}(x(t) - y(t))$ is

$$\beta\,T = \bigcup_{n\in\mathbb{Z}}\left(\arctan(\theta) + n\,\pi,\ \frac{\pi}{2} + n\,\pi\right), \qquad \theta = \frac{\beta^2 - \gamma\,\delta^2}{\beta\,\delta\,(1+\gamma)}.$$

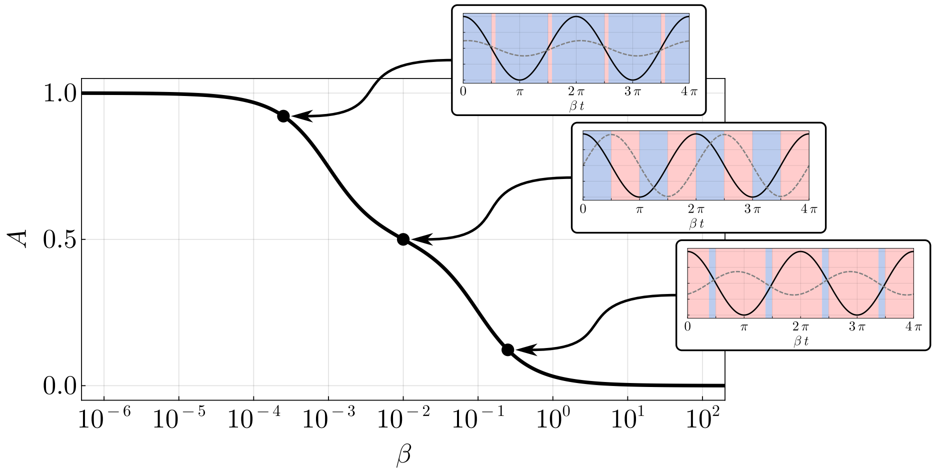


Figure 3. Prediction accuracy over an interval giving an oscillating input (14). In the low-frequency limit ($\beta \to 0^+$), the system achieves perfect predictive performance, with accuracy $A \to 1$. Conversely, in the high-frequency regime ($\beta \to \infty$), the prediction mechanism fails systematically, $A \to 0$. Parameters were fixed at $\alpha = 10^{-1}$, $\gamma = 10^{-2}$, and $\delta = 10^{-1}$.

After each sign change of the derivative, there is a delay $\arctan(\theta)$ where predictions still follow the previous trend.

In the limiting regime where $\beta \to 0^+$, $\arctan(\theta) \to -\pi/2$, which approximates the expression for the derivative of the input (14). Consequently, the temporal delay between a sign change in the derivative and the subsequent prediction sign change becomes small compared to input dynamics. This fast response of the prediction to the derivative's trend indicates a state of near in-phase synchronization between the predicted and the real trend.

Conversely, as $\beta \to \infty$, $\arctan(\theta) \to \pi/2$. Under this condition, a significant delay emerges between a sign change in the derivative and the interval where the prediction accurately reflects the trend. The prediction tends to maintain its sign for a long time after a change in the derivative's sign. This delayed response results in the predicted sign evolving in a manner nearly opposite to the input's trend, with a near anti-phase behavior between the predicted and the real trend.

The prediction accuracy $A$ is defined as the proportion of time the system produces correct predictions within a given period. This is obtained by calculating the ratio between the union of all correct prediction intervals and the total period length. Given that these correct intervals exhibit periodicity with a spacing of $\pi$ in the scaled time domain $\beta\, t$, the normalized accuracy reduces to

$$A = \frac{1}{2} - \frac{1}{\pi} \arctan(\theta),$$

which ranges over $[0, 1]$—corresponding to completely misaligned ($A = 0$) and perfectly aligned ($A = 1$) cases. This follows from the fact that

$$-\frac{\pi}{2} < \arctan(\theta) < \frac{\pi}{2}, \qquad \theta \in \mathbb{R}.$$

Figure 3 displays the prediction accuracy $A$ across a relevant frequency range. As shown, the system remains nearly in phase with the input trend under slow dynamics. In contrast, faster dynamics drive the system toward an anti-phase response.

It is noteworthy to notice that while predictions are nearly correct over an entire period as $\beta \to 0^+$ and almost entirely wrong as $\beta \to \infty$, the magnitude $|x - y|$ vanishes. This behavior is further investigated when additive noise is introduced into the system (see Section II F).

### II. STOCHASTIC DYNAMICS

#### A. Wiener Process Predictability

Let's start by setting up a basic probability framework for stochastic processes [2]. A probability space is formally represented by the triplet $(\Omega,\ \mathcal{F},\ \mathbb{P})$, where:

- $\Omega$ denotes the sample space, the set of all possible elementary outcomes $\omega$ of a random experiment.
- $\mathcal{F}$ is a $\sigma$-algebra on $\Omega$, which is a collection of subsets of $\Omega$ satisfying three axioms:
  - $\Omega \in \mathcal{F}$.
  - Closed under complementation. If $A \in \mathcal{F}$, then $A^c \in \mathcal{F}$.
  - Closed under countable unions. If $A_1, A_2, \dots \in \mathcal{F}$, then $\bigcup_{i=1}^{\infty} A_i \in \mathcal{F}$s.

  Elements of $\mathcal{F}$ are termed events, representing measurable occurrences to which probabilities can be assigned.
- $\mathbb{P} : \mathcal{F} \to [0,1]$ is the probability measure, a function satisfying Kolmogorov's axioms: non-negativity, $\mathbb{P}(\Omega) = 1$, and countable additivity for disjoint events.

A stochastic process $\{u(t)\}_{t\geq t_0}$ is a family of random variables indexed by a time parameter $t$, where each $u(t) : \Omega \to \mathbb{R}$ is a $\mathcal{F}$-measurable function [3]. This measurability ensures that for any Borel set $B \subseteq \mathbb{R}$, the preimage

$$u(t)^{-1}(B) = \{\omega \in \Omega : u(t)(\omega) \in B\}$$

belongs to $\mathcal{F}$, thereby permitting the assignment of probabilities to events involving $u(t)$. The process evolves over time, with $u(t)$ capturing the state of the system at time $t$.

Focus on stochastic processes given by $u(t) \coloneqq W(t)$, where $W(t)$ represents a standard Wiener process, defined by its characteristic properties:

1. $W(t_0) = 0$ almost surely, meaning $\mathbb{P}(W(t_0) = 0) = 1$.
2. For any finite sequence $t_0 \leq t_1 < t_2 < \cdots < t_n$, the increments $W(t_{k+1}) - W(t_k)$ are mutually independent random variables.
3. For $t_i < t_j$, the increment $W(t_j) - W(t_i)$ follows a normal distribution

   $$W(t_j) - W(t_i) \sim \mathcal{N}(0,\ t_j - t_i).$$

   This implies that the dispersion of increments scales with the square root of the time interval, a hallmark of diffusive behavior.

Central to the analysis of stochastic processes is the natural filtration $\{\mathcal{F}(t)\}_{t\geq t_0}$, where $\mathcal{F}(t)$ is the smallest $\sigma$-algebra generated by the history of the process up to time $t$. This filtration encapsulates all available information about the trajectory of $W(t)$ until time $t$.

For the Wiener process, the conditional probability of observing an upward trend in an infinitesimal interval $[t,\ t+\delta\, t)$ satisfies

$$\mathbb{P}\left(\Delta W(t) > 0 \mid \mathcal{F}(t)\right) = \frac{1}{2},$$

where $\Delta W(t) = W(t + \delta\, t) - W(t)$. This equality arises from two fundamental properties:

- Independent Increments: The future increment $\Delta W(t)$ is statistically independent of $\mathcal{F}(t)$.
- Symmetric Distribution: The increment $\Delta W(t)$ follows a normal distribution with zero mean, which is symmetric with respect to the origin.

As a consequence, the likelihood of an upward movement is identical to that of a downward movement at any given time, regardless of the process's past trajectory. Thus, attempting to predict the trend of a Wiener process cannot be expected to be more effective than random guessing.

This lack of directional bias is also demonstrated by the conditional expectation of the increments

$$\mathbb{E}\left[\Delta W(t) \mid \mathcal{F}(t)\right] = 0.$$

The expectation vanishes because the distribution of $\Delta W(t)$ is symmetric, and increments with the same magnitude and different sign cancel out in expectation. This result also shows that the Wiener process is a martingale with respect to its natural filtration, embodying the "efficient market hypothesis" in financial mathematics: past movements provide no exploitable information for predicting future behavior.

#### B. Probability of a Random Guesser Correctly Predicting a Trend

Consider a random binary guesser attempting to predict the trend $\Delta u(t)$ of a random input signal $u(t)$. The trend $\Delta u(t)$ can either be positive or negative, and the guesser has no prior knowledge of the underlying process generating. Thus, the guesser's prediction is correct with probability $p = 1/2$ at each trial, independent of past predictions and any biases in $u$.

Let $X_i$ be a Bernoulli random variable representing the outcome of the $i$-th prediction:

$$X_i = \begin{cases} +1 & \text{with probability } p = \frac{1}{2}, \\ -1 & \text{with probability } 1 - p = \frac{1}{2}. \end{cases}$$

For $M$ independent trials, the total number of correct predictions $K_M$ is

$$K_M = \sum_{i=1}^{M} X_i.$$

Since the $X_i$ are i.i.d. Bernoulli trials, $K_M$ follows a binomial distribution:

$$K_M \sim \text{Binomial}\left(M,\ \frac{1}{2}\right).$$

The probability mass function is

$$\mathbb{P}(K_M = k) = \binom{M}{k} \left(\frac{1}{2}\right)^k \left(\frac{1}{2}\right)^{M-k} = \binom{M}{k} \left(\frac{1}{2}\right)^M,$$

where $\binom{M}{k}$ is the binomial coefficient.

The cummulative distribution function (CDF) $F_{K_M}(k)$ gives the probability that the number of correct predictions is at most $k$:

$$F_{K_M}(k) = P(K_M \leq k) = \sum_{i=0}^{k} \binom{M}{i} \left(\frac{1}{2}\right)^M,$$

and its complement survival function

$$S_{K_M}(k) = P(K_M > k) = 1 - F_{K_M}(k).$$

For $k = \frac{M}{2}$, the CCDF represents the probability that the guesser performs strictly better than random chance (i.e., more than half predictions are correct).

For large $M$, the binomial distribution can be approximated using the *De Moivre-Laplace theorem*, a special case of the Central Limit Theorem (CLT) for binomial variables [4]. The theorem states that for large number of samples $M$, the standardized binomial distribution approaches the standard normal distribution $\mathcal{N}(0,\ 1)$:

$$\frac{K_M - \mu_n}{\sigma_n} \xrightarrow{d} \mathcal{N}(0, 1),$$

with mean $\mu_M = M\,p$, and variance $\sigma_M^2 = M\,p\,(1-p)$.

Thus, for large $M$,

$$\lim_{M\to\infty} K_M \approx \mathcal{N}\left(\frac{M}{2},\ \frac{M}{4}\right).$$

Let $\Phi(z)$ denote the standard normal CDF. Then the survival function for $K_M > \frac{M}{2}$ can be expressed as

$$\mathbb{P}\left(K_M > \frac{M}{2}\right) = 1 - \mathbb{P}\left(K_M \leq \frac{M}{2}\right) \approx 1 - \Phi\left(\frac{\frac{M}{2} - \frac{M}{2}}{\frac{\sqrt{M}}{2}}\right) = 1 - \Phi(0),$$

and since $\Phi(0) = \frac{1}{2}$,

$$\mathbb{P}\left(K_M > \frac{M}{2}\right) \approx 1 - \frac{1}{2} = \frac{1}{2}.$$

#### C. Probability of a Random Guesser Outperforming Another

Consider two independent agents, $G_1$ and $G_2$, each making a sequence of $N$ binary predictions where the true outcome at each step is equally likely to be either value. Both guessers have no predictive information, meaning each of their predictions is correct with probability $\frac{1}{2}$, independently of all other predictions. The probability that, after $N$ trials, $G_1$ achieves strictly more correct predictions than $G_2$, denoted $\mathbb{P}(c_1 > c_2)$, where $c_1$ and $c_2$ are the total numbers of correct predictions for each guesser, respectively.

Since both guessers are statistically identical and independent, their correct prediction counts, $c_1$ and $c_2$, follow identical binomial distributions:

$$c_1, c_2 \sim \text{Binomial}\left(N,\ \frac{1}{2}\right).$$

By symmetry, the probability that $G_1$ outperforms $G_2$ is equal to the probability that $G_2$ outperforms $G_1$:

$$\mathbb{P}(c_1 > c_2) = \mathbb{P}(c_2 > c_1).$$

The remaining probability corresponds to the event where both guessers have the same number of correct predictions, $\mathbb{P}(c_1 = c_2)$. These probabilities must sum to 1, leading to the relationship $2\mathbb{P}(c_1 > c_2) + \mathbb{P}(c_1 = c_2) = 1$, which can be rearranged to express $\mathbb{P}(c_1 > c_2)$ in terms of $\mathbb{P}(c_1 = c_2)$:

$$\mathbb{P}(c_1 > c_2) = \frac{1 - \mathbb{P}(c_1 = c_2)}{2}.$$

The key challenge, therefore, reduces to calculating $\mathbb{P}(c_1 = c_2)$. Because $c_1$ and $c_2$ are independent, this probability is given by the sum over all possible values $k$ of the product of their individual probabilities:

$$\mathbb{P}(c_1 = c_2) = \sum_{k=0}^{N} \mathbb{P}(c_1 = k)\, \mathbb{P}(c_2 = k) = \sum_{k=0}^{N} \binom{N}{k}^2 \left(\frac{1}{2}\right)^{2N}.$$

This sum can be simplified using a combinatorial identity. Specifically, the sum of squared binomial coefficients $\sum_{k=0}^{N} \binom{N}{k}^2$ is equal to the central binomial coefficient $\binom{2N}{N}$. Thus, the probability of a tie simplifies to

$$\mathbb{P}(c_1 = c_2) = \frac{1}{4^N} \binom{2N}{N}.$$

Substituting this back into the expression for $\mathbb{P}(c_1 > c_2)$, the probability becomes

$$\mathbb{P}(c_1 > c_2) = \frac{1}{2}\left(1 - \frac{1}{4^N}\binom{2N}{N}\right).$$

For large $N$, the central binomial coefficient $\binom{2N}{N}$ can be approximated using Stirling's formula, yielding

$$\binom{2N}{N} \approx \frac{4^N}{\sqrt{\pi N}}.$$

Consequently, $\mathbb{P}(c_1 = c_2)$ behaves asymptotically as $\sqrt{\pi N}^{-1}$, which tends to 0 as $N \to \infty$. This implies that the probability of a tie becomes negligible for large $N$, and $\mathbb{P}(c_1 > c_2)$ converges to $\frac{1}{2}$.

Thus, as the number of trials increases, the chance that one guesser outperforms the other approaches 1/2, with the vanishing probability mass corresponding to the increasingly rare event of an exact tie.

#### D. General Drift Input with Additive Noise

Given a general input signal $u(t)$ in the stochastic system described by the Ornstein–Uhlenbeck (OU) processes:

$$\begin{aligned} dx &= [\alpha\, u(t) - \delta\, x]\; dt + \eta\, \alpha\, dW_x(t), \\ dy &= \gamma\; [\alpha\, u(t) - \delta\, y]\; dt + \eta\, \gamma\, \alpha\, dW_y(t), \end{aligned} \tag{17}$$

a stochastic extension of system (1), where the additive noise terms model environmental fluctuations proportional the sensor constants that perturb the system's deterministic dynamics. Parameter $\eta$ scales the noise intensity and $dW(t)$ represents independent increments of a Wiener process (Brownian motion), capturing unpredictable, continuous-time random disturbances.

This is a linear system of stochastic differential equations (SDEs) of the form

$$dy = (a(t)\, y + b(t))\; dt + c(t)\, dW(t),$$

where $a(t) \coloneqq -\gamma\, \delta$, $b(t) \coloneqq \gamma\, \alpha\, u(t)$, and $c(t) \coloneqq \eta\, \gamma\, \alpha$. This can be solved by integrating factors as the deterministic case. The general solution for such linear SDE is

$$y(t) = e^{A(t)} \left( y(0) + \int_0^t e^{-A(s)}\, b(s)\, ds + \int_0^t e^{-A(s)}\, c(s)\, dW(s) \right),$$

where $A(t) = \int_0^t a(s)\, ds = -\gamma\, \delta\, t$.

The first integral (drift term) is

$$\int_0^t e^{\gamma\, \delta\, s}\, b(s)\, ds = \gamma\, \alpha \int_0^t e^{\gamma\, \delta\, s}\, u(s)\, ds,$$

and the second integral (diffusion term) is

$$\int_0^t e^{\gamma\, \delta\, s}\, c(s)\, dW(s) = \eta\, \gamma\, \alpha \int_0^t e^{\gamma\, \delta\, s}\, dW(s),$$

which allows to express the solution as

$$y(t) = e^{-\gamma\, \delta\, t} \left( y_0 + \gamma\, \alpha \int_0^t e^{\gamma\, \delta\, s}\, u(s)\, ds + \eta\, \gamma\, \alpha \int_0^t e^{\gamma\, \delta\, s}\, dW(s) \right).$$

As the dynamics of $x(t)$ are equivalent to $y(t)$ when $\gamma = 1$, the solution is given by

$$x(t) = e^{-\delta\, t} \left( x_0 + \alpha \int_0^t e^{\delta\, s}\, u(s)\, ds + \eta\, \alpha \int_0^t e^{\delta\, s}\, dW(s) \right).$$

The fact that both $x$ and $y$ are Gaussian OU processes implies that they are completely specified by their first two statistical moments.

The first moment $\mathbb{E}[x(t)]$ and $\mathbb{E}[y(t)]$ follows the deterministic part of the OU processes:

$$\mathbb{E}[x(t)] = x_0\, e^{-\delta\, t} + \alpha \int_0^t u(s)\, e^{-\delta\,(t-s)}\, ds, \quad \mathbb{E}[y(t)] = y_0\, e^{-\gamma\, \delta\, t} + \gamma\, \alpha \int_0^t u(s)\, e^{-\gamma\, \delta\,(t-s)}\, ds. \tag{18}$$

Thus, the expected difference of moving averages is

$$\mu(t) \coloneqq \mathbb{E}[x(t) - y(t)] = \mathbb{E}[x(t)] - \mathbb{E}[y(t)],$$

which at long times can be approximated as

$$\mathbb{E}[x(t) - y(t)] \approx \alpha \int_0^t u(s) \left( e^{-\delta\,(t-s)} - \gamma\, e^{-\gamma\, \delta\,(t-s)} \right) ds.$$

If $u(t)$ is time-varying, $\mu(t)$ is a convolution of $u(t)$ with the impulse response difference. If the input is constant, $u(t) = u_0$, the long-term expected difference converges to zero.

The second moment, $\mathrm{Var}[x(t)]$ and $\mathrm{Var}[y(t)]$ is independent of the drift as it is deterministically defined and affects only the expectation.

To compute variances of stochastic processes, the Itô isometry is usually used to simplify the computations. Given an adapted stochastic process $g(t)$ such that $\mathbb{E}\left[\int_0^t g(t)^2\, ds\right] < \infty$, the Itô isometry states:

$$\mathbb{E}\left[\left(\int_0^t g(s)\, dW(s)\right)^2\right] = \mathbb{E}\left[\int_0^t g(s)^2\, ds\right]. \tag{19}$$

The variance of $y(t)$ comes entirely from the stochastic integral term (since the other terms are deterministic). Thus:

$$\mathrm{Var}[y(t)] = \mathbb{E}\left[\left(e^{-\gamma\,\delta\,t}\,\eta\,\gamma\,\alpha \int_0^t e^{\gamma\,\delta\,s}\, dW(s)\right)^2\right] = e^{-\gamma\,\delta\,t}\,\eta\,\gamma\,\alpha\;\mathbb{E}\left[\left(\int_0^t e^{\gamma\,\delta\,s}\, dW(s)\right)^2\right].$$

Using the Itô isometry (19):

$$\mathbb{E}\left[\left(\int_0^t e^{\gamma\,\delta\,s}\, dW(s)\right)^2\right] = \mathbb{E}\left[\int_0^t e^{2\,\gamma\,\delta\,s}\, ds\right],$$

which can be computed as

$$\int_0^t e^{2\gamma\,\delta s}\, ds = \left.\frac{e^{2\gamma\,\delta s}}{2\gamma\,\delta}\right|_0^t = \frac{e^{2\gamma\,\delta\,t} - 1}{2\gamma\,\delta}.$$

Substitute back into the variance expression:

$$\mathrm{Var}[y(t)] = \left(e^{-\gamma\,\delta\,t}\,\eta\,\gamma\,\alpha\right)^2\,\frac{e^{2\,\gamma\,\delta\,t} - 1}{2\,\gamma\,\delta} = \frac{\eta^2\,\gamma\,\alpha^2}{2\,\delta}\,\left(1 - e^{-2\,\gamma\,\delta\,t}\right).$$

Since the dynamics of $x(t)$ are equivalent to $y(t)$ when $\gamma = 1$, its variance is given by

$$\mathrm{Var}[x(t)] = \frac{\eta^2\,\alpha^2}{2\,\delta}\left(1 - e^{-2\,\delta\,t}\right).$$

Assuming correlated noise, $dW := dW_x = dW_y$, as both variables are driven by the same input, the variance of the difference of moving averages becomes

$$\sigma^2(t) := \mathrm{Var}[x(t) - y(t)] = \mathrm{Var}[x(t)] + \mathrm{Var}[y(t)] - 2\mathrm{Cov}[x(t), y(t)].$$

Given both variables have zero expectation, the covariance is

$$\mathrm{Cov}[x(t),\ y(t)] = \mathbb{E}\left[x(t)\,y(t)\right] = \mathbb{E}\left[\left(\eta\,\alpha \int_0^t e^{-\delta\,(t-s)}\, dW(s)\right)\left(\eta\,\gamma\,\alpha \int_0^t e^{-\gamma\,\delta\,(t-s)}\, dW(s)\right)\right],$$

which can be solved using Itô isometry (19):

$$\mathrm{Cov}[x(t),\ y(t)] = \eta^2\,\gamma\,\alpha^2 \int_0^t e^{-\delta\,(t-s)}\,e^{-\gamma\,\delta\,(t-s)}\, ds = \frac{\eta^2\,\gamma\,\alpha^2}{\delta\,(1+\gamma)}\left(1 - e^{-\delta\,(1+\gamma)\,t}\right).$$

Substituting the variances and covariance,

$$\sigma^2(t) = \frac{\eta^2\,\alpha^2}{2\,\delta}\left(1 - e^{-2\,\delta\,t}\right) + \frac{\eta^2\,\gamma\,\alpha^2}{2\,\delta}\left(1 - e^{-2\,\gamma\,\delta\,t}\right) - \frac{2\,\eta^2\,\gamma\,\alpha^2}{\delta\,(1+\gamma)}\left(1 - e^{-\delta\,(1+\gamma)\,t}\right).$$

In the long-term stationary limit ($t \to \infty$),

$$\sigma_\infty^2 \coloneqq \lim_{t\to\infty} \sigma^2(t) = \frac{\eta^2\,\alpha^2\,(1-\gamma)^2}{2\,\delta\,(1+\gamma)}.$$

Thus, for large $t$, difference $x(t) - y(t)$ is approximately normally distributed,

$$x(t) - y(t) \sim \mathcal{N}(\mu(t),\ \sigma_\infty^2),$$

with a stationary variance that does not depend on $u(t)$, owing to the additive nature of the noise.

In the case where $\gamma = 1$, mean and variance for the difference of moving averages vanish ($\mu(t) = 0$ and $\sigma(t)^2 = 0$), as both variables follow the same deterministic dynamics and noise is completely correlated.

### E. Constant Drift Input with Additive Noise

In the specific case when $u(t) = \beta\,t + u_0$, the expected values $\mathbb{E}[x(t)]$ and $\mathbb{E}[y(t)]$ can be computed explicitly as integrals following (18):

$$\begin{aligned}
\mathbb{E}[x(t)] &= x_0\,e^{-\delta\,t} + \alpha \int_0^t (\beta\,s + u_0)\; e^{-\delta\,(t-s)}\,ds,\\
\mathbb{E}[y(t)] &= y_0\,e^{-\gamma\,\delta\,t} + \gamma\,\alpha \int_0^t (\beta\,s + u_0)\; e^{-\gamma\,\delta\,(t-s)}\,ds,
\end{aligned}$$

which are the same expressions as in the deterministic case (6). The variances remain unchanged from the general case, as they do not depend on $u(t)$.

The expectation $\mu \coloneqq \mathbb{E}[x(t) - y(t)]$ and the variance $\sigma^2 \coloneqq \mathrm{Var}[x(t) - y(t)]$ are

$$\begin{aligned}
\mu(t) &= \frac{\alpha\,\beta}{\delta^2\,\gamma}\,(1-\gamma) + \left(\frac{\alpha\,\beta}{\delta^2} - \frac{\alpha\,u_0}{\delta} + x_0\right) e^{-\delta\,t} - \left(\frac{\alpha\,\beta}{\delta^2\,\gamma} - \frac{\alpha\,u_0}{\delta} + y_0\right) e^{-\delta\,\gamma\,t},\\
\sigma^2(t) &= \frac{\eta^2\,\alpha^2}{2\,\delta}\left(1 - e^{-2\,\delta\,t}\right) + \frac{\eta^2\,\gamma\,\alpha^2}{2\,\delta}\left(1 - e^{-2\,\gamma\,\delta\,t}\right) - \frac{2\,\eta^2\,\gamma\,\alpha^2}{\delta\,(1+\gamma)}\left(1 - e^{-\delta\,(1+\gamma)\,t}\right).
\end{aligned}$$

Thus, at long times, the process converges as a Gaussian distribution with finite mean and variance:

$$\mu_\infty = \lim_{t\to\infty} \mu(t) = \frac{\alpha\,\beta}{\delta^2\,\gamma}\,(1-\gamma)\,, \qquad \sigma_\infty^2 = \lim_{t\to\infty} \sigma^2(t) = \frac{\eta^2\,\alpha^2\,(1-\gamma)^2}{2\,\delta\,(1+\gamma)},$$

with both values vanishing when $\gamma = 1$.

#### *Probability of Correct Predictions with Noise*

The expected trend, given by the sign of the mean $\mathrm{sgn}(\mu(t))$ correctly predicts the trend, but due to the variance, individual trajectories might incorrectly predict at any point in time. Consider the actual value at any time as a random variable $z(t) \coloneqq \mu(t) + \mathcal{N}\left(0,\ \sigma(t)^2\right)$.

- If $\mu > 0$,

$$\mathbb{P}(z > 0) = \mathbb{P}(\mu + N > 0) = \mathbb{P}(N > -\mu) = \mathbb{P}\left(\frac{N}{\sigma} > -\frac{\mu}{\sigma}\right) = 1 - \mathbb{P}\left(\frac{N}{\sigma} \le -\frac{\mu}{\sigma}\right)) = 1 - \Phi\left(-\frac{\mu}{\sigma}\right) = \Phi\left(\frac{\mu}{\sigma}\right),$$

  using the property $\Phi(-z) = 1 - \Phi(z)$.

- If $\mu < 0$,

$$\mathbb{P}(X < 0) = \mathbb{P}(\mu + N < 0) = \mathbb{P}(N < -\mu) = \mathbb{P}\left(\frac{N}{\sigma} < -\frac{\mu}{\sigma}\right) = \Phi\left(-\frac{\mu}{\sigma}\right).$$

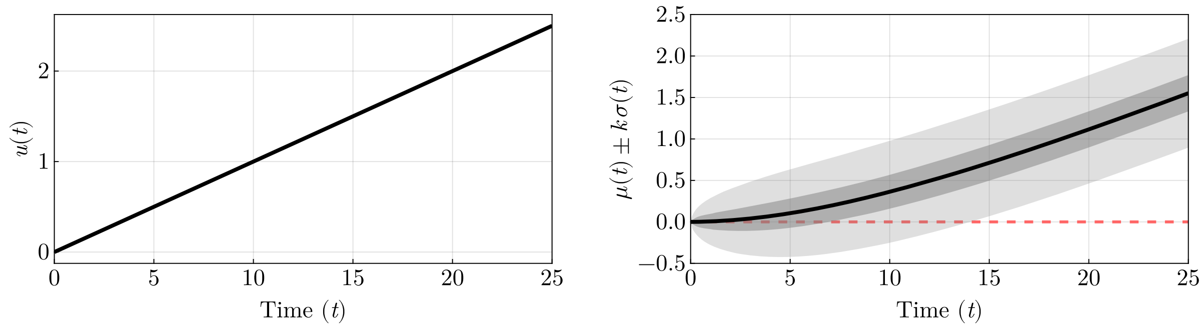


Figure 4. Left panel shows a time-series of an input with constant drift. Right panel displays the expected system prediction $\mu(t) = E[x(t) - y(t)]$, with uncertainty bands showing $\pm\sigma$ ($\sim$68 % confidence, dark gray shadow) and $\pm 3\,\sigma$ ($\sim$99.7 % confidence, light gray shadow). The variance $\sigma^2$ captures transient and steady-state stochasticity, with bands initially tight (zero uncertainty at $t = 0$) and asymptotically approaching constant width as the system reaches statistical equilibrium. As it can be seen, as the input increases in concentration, almost all trajectories will successfully predict the correct trend. Parameters were fixed at $\alpha = 10^{-1}$, $\gamma = 10^{-2}$, $\delta = 10^{-1}$, $\beta = 10^{-1}$, and $\eta = 1$.

Thus given that $\mathrm{sgn}(\mu(t)) = \mathrm{sgn}(\beta)$, the probability of a correct prediction is given by $\mathbb{P}\left(|\mu(t)| > 0\right)$, due to the symmetry of the distribution:

$$\mathbb{P}\left(|\mu(t)| > 0\right) = \Phi\left(\frac{|\mu|}{\sigma}\right).$$

As shown in Figure 4, during the recovery phase following a drift change, most trajectories align with the sign of the deterministic mean—provided the system is not noise-dominated, i.e., $|\mu_\infty| \gg \sigma_\infty$.

#### F. Oscillating Input with Additive Noise

In the specific case when $u(t) = \sin(\beta\, t) + u_0$, the expected values $\mathbb{E}[x(t)]$ and $\mathbb{E}[y(t)]$ can be computed explicitly as integrals following (18):

$$\mathbb{E}[x(t)] = x_0\, e^{-\delta\, t} + \alpha \int_0^t \left(\sin(\beta\, s) + u_0\right)\, e^{-\delta\,(t-s)}\, ds,$$
$$\mathbb{E}[y(t)] = y_0\, e^{-\gamma\,\delta\, t} + \gamma\,\alpha \int_0^t \left(\sin(\beta\, s) + u_0\right)\, e^{-\gamma\,\delta\,(t-s)}\, ds,$$

which are the same expressions as in the deterministic case (15). The variances remain unchanged from the general case, as they do not depend on $u(t)$.

Let $\mu(t) := \mathbb{E}[x(t) - y(t)]$ denote the mean and $\sigma^2(t) := \mathrm{Var}[x(t) - y(t)]$ the variance of the difference process. Asymptotically, this process converges in distribution to a Gaussian whose mean evolves in time and settles into a steady-state oscillation. This limiting mean can be expressed in amplitude–phase form as

$$\mu(t) \to \sqrt{C^2 + S^2}\, \cos\left(\beta\, t + \arctan\left(\frac{S}{C}\right)\right),$$

where the constants $C$ and $S$ are given by

$$C := \left(\frac{\alpha\,\beta\gamma}{\beta^2 + \delta^2\,\gamma^2} - \frac{\alpha\,\beta}{\beta^2 + \delta^2}\right), \qquad S := \left(\frac{\alpha\delta\gamma^2}{\beta^2 + \delta^2\,\gamma^2} - \frac{\alpha\delta}{\beta^2 + \delta^2}\right),$$

and the constant stationary variance, independent of $u(t)$, is given by

$$\sigma_\infty^2 = \frac{\eta^2\,\alpha^2\,(1-\gamma)^2}{2\,\delta\,(1+\gamma)}.$$

Following the deterministic dynamics, the expectation is correct in the intervals

$$\beta T = \bigcup_{n\in\mathbb{Z}} \left(\arctan(\theta) + n\,\pi,\ \frac{\pi}{2} + n\,\pi\right), \qquad \theta = \frac{\beta^2 - \gamma\,\delta^2}{\beta\,\delta\,(1+\gamma)}.$$

However, the introduction of noise causes some trajectories within these intervals of correct expectation to deviate and incorrectly predict the trend. Conversely, noise can also lead trajectories in the complementary intervals where the deterministic expectation is wrong to correctly predict the underlying trend.

This interplay between deterministic behavior and noise becomes particularly significant in the limiting cases when $\beta \to 0$ and $\beta \to \infty$.

When the input changes slowly ($\beta \to 0$), one would expect the system to easily track the trend, leading to almost always correct predictions as the system can rapidly adapt to the sign changes. However, the small magnitude of the difference $|x - y|$ in this regime means the stochastic terms become dominant, reducing the probability of a correct prediction to random.

Conversely, when the input changes rapidly ($\beta \to \infty$), the expectation is that predictions will be out of phase with the input, leading to incorrect predictions most of the time. Yet again, due to the small value of $|x-y|$, the stochastic terms take over, counterintuitively increasing the probability of a correct prediction to random.

*Probability of Correct Predictions with Noise*

An analytical derivation of the correct prediction probability involves considering two types of intervals: those where the expected trend is correct, but noise can lead to incorrect predictions, and those where the expected trend is incorrect, yet noise can result in correct predictions.

When the expected prediction aligns with the true trend, $\mathrm{sgn}(\mu(t)) = \mathrm{sgn}(\cos(\beta t))$, the system exhibits a natural bias toward correct predictions. In this regime, the probability of maintaining the correct sign despite noise perturbations is given by the upper tail of the noise distribution. This corresponds to

$$\mathbb{P}(t) = \Phi\left(\frac{|\mu(t)|}{\sigma}\right),$$

where $\Phi(\cdot)$ denotes the cumulative distribution function of the standard normal distribution. This expression gives the probability that a zero-mean Gaussian noise term with variance $\sigma^2$ does not exceed $|\mu(t)|$ in magnitude, thereby preserving the sign of the deterministic trend prediction $\mu(t)$.

Conversely, when the mean prediction opposes the true trend, $\mathrm{sgn}(\mu(t)) \neq \mathrm{sgn}(\cos(\beta t))$, the system is inherently biased toward incorrect predictions. In this scenario, a correct prediction requires the noise to be sufficiently strong to flip the sign of the observed difference. The probability of this occurring is quantified by the lower tail of the noise distribution, expressed as

$$\mathbb{P}(t) = \Phi\left(-\frac{|\mu(t)|}{\sigma}\right).$$

This reflects the likelihood that the noise term overcomes the misleading expectation $\mu(t)$ to produce an observation consistent with the true trend.

The time-dependent probability $\mathbb{P}(t)$ thus alternates between these two cases:

$$\mathbb{P}(t) = \begin{cases} \Phi\left(+\frac{|\mu(t)|}{\sigma}\right) & \text{if } \mathrm{sgn}(\mu(t)) = \mathrm{sgn}(\cos(\beta t)), \\ \Phi\left(-\frac{|\mu(t)|}{\sigma}\right) & \text{if } \mathrm{sgn}(\mu(t)) \neq \mathrm{sgn}(\cos(\beta t)). \end{cases}$$

As shown in Figure 5, the prediction probability drops abruptly at each input trend switch, after which most trajectories recover and gradually increase the probability of a correct prediction until the next switch.

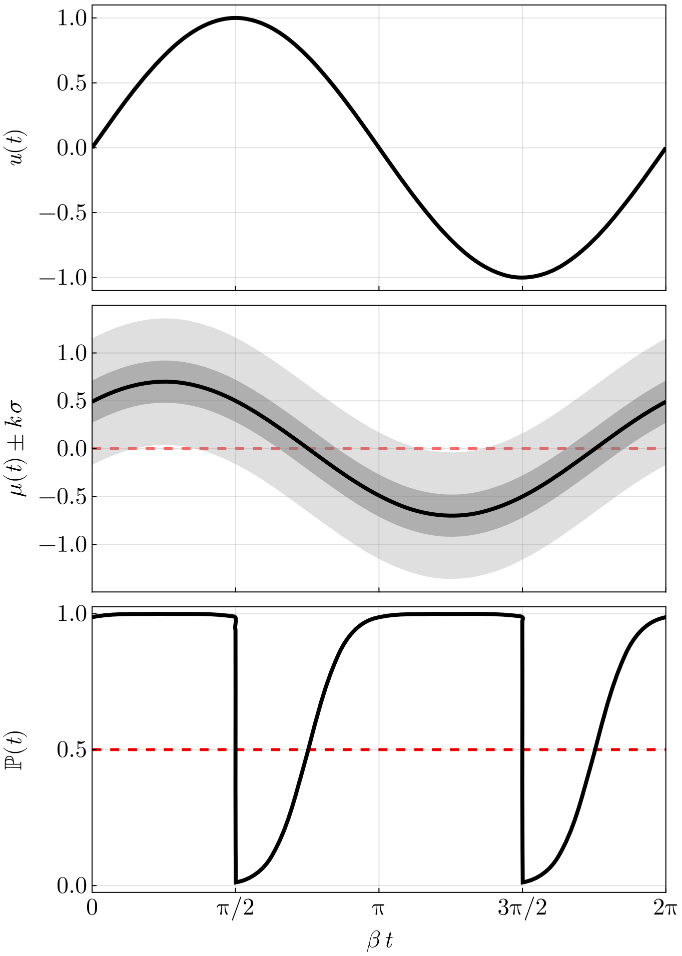


Figure 5. Top panel shows a time-series of an oscillating input $u(t)$. Middle panel displays the expected system prediction $\mu(t) = E[x(t) - y(t)]$, with uncertainty bands showing $\pm\sigma$ ($\sim$68 % confidence, dark gray shadow) and $\pm 3\,\sigma$ ($\sim$99.7 % confidence, light gray shadow). Bottom panel presents the time-dependent probability $\mathbb{P}(t)$ of correct trend prediction, relative to a random guesser (red dashed line). Parameters were fixed at $\alpha = 10^{-1}$, $\gamma = 10^{-2}$, $\delta = 10^{-1}$, $\beta = 10^{-3}$, and $\eta = 1$.

### III. CODE AVAILABILITY

The code used for the simulations is available under a permissive license [5].
